# Genotoxic and metabolic stress drive divergent senescence programs in human microglia

**DOI:** 10.64898/2026.08.31.748228

**Authors:** Nisha V. Jose, Susan Janssens, Eunchai Kang, Bettina Platt

**Affiliations:** School of Medicine, Medical Sciences and Nutrition, Institute of Medical Sciences, University of Aberdeen, Aberdeen, AB25 2ZD, United Kingdom

**Keywords:** Microglial senescence, Human Microglial Cell Line, Genotoxic stress, Metabolic stress, Mitochondrial dysfunction, Neuroinflammation

## Abstract

Microglial dysfunction is a hallmark of brain ageing linked to the accumulation of senescent microglial phenotypes that promote chronic neuroinflammation. Both genotoxic and metabolic stress have been implicated in microglial senescence; yet, whether distinct stressors shape senescence programs remain unclear. Here, we investigated the impact of chronic genotoxic and metabolic stress on senescence-associated phenotypes in the human microglia cell line HMC3. Cells were exposed to doxorubicin to induce sustained DNA damage or to chronic high-glucose conditions to model metabolic stress. Both stress paradigms induced characteristic senescence features including cellular and nuclear hypertrophy, increased senescence-associated β-galactosidase activity and reduced metabolic viability without significant cell loss. Both conditions activated the p53-p21 pathway and sustained DNA damage signalling, whereas metabolic stress additionally induced p16 expression and peripheral nuclear localisation of p21, suggesting divergence in senescence regulatory pathways. Mitochondrial alterations were evident under both conditions, Dox-induced stress was associated with downregulation of NRF2-TFAM signalling, whereas HG-induced stress induced NRF2-TFAM activation alongside increased KEAP1 expression, suggesting a constrained antioxidant response. This was accompanied by activation of mitochondrial and antioxidant stress responses that did not restore mitochondrial content. Furthermore, both stressors induced robust inflammatory activation, with genotoxic stress promoting a chemokine-rich senescence-associated secretory phenotype, while metabolic stress induced an interferon-associated inflammatory signature. Collectively, these findings demonstrated that chronic genotoxic and metabolic stress drive distinct yet overlapping senescence programs characterised by morphological changes, mitochondrial remodelling and persistent inflammatory activation. These stress-specific responses may differentially contribute to neurodegenerative processes and disease susceptibility.

## Introduction

Ageing is the primary risk factor for neurodegenerative diseases and is associated with profound alterations in brain immune homeostasis, with microglia undergoing functional, metabolic and transcriptional reprogramming that collectively contribute to chronic neuroinflammation. In the aged brain, microglia shift from a homeostatic surveillance state towards chronically inflammatory phenotype, characterised by impaired phagocytosis, altered metabolic flexibility and enhanced inflammatory signalling [1,2]. These age-associated microglial changes are strongly implicated in the progression of neurodegenerative disorders, including Alzheimer’s disease, where persistent microglial activation exacerbates synaptic dysfunction and neuronal vulnerability [3,4].

An emerging contributor to age-related microglial dysfunction is cellular senescence. Senescent cells are defined by stable cell-cycle arrest accompanied by distinct morphological changes, chromatin remodelling, lysosomal expansion, mitochondrial dysfunction and the development of a senescence-associated secretory phenotype (SASP) [5,6]. While originally described in proliferative cells, accumulating evidence indicates that post-mitotic or slowly dividing cells, including microglia, can adopt senescence-like phenotypes in response to chronic stress [7]. In aged and diseased brains, these senescent microglia promote chronic neuroinflammation through sustained secretion of pro-inflammatory mediators rather than undergoing apoptotic clearance [8,9], ultimately exacerbating disease progression via persistent inflammatory signalling, impaired clearance of cellular debris and amyloid aggregates, and disruption of metabolic homeostasis [10,11].

Multiple stressors have been proposed to drive microglial senescence, with genotoxic and metabolic stress emerging as particularly relevant. Persistent DNA damage accumulates in ageing microglia as a consequence of oxidative stress, impaired DNA repair and repeated inflammatory activation, leading to activation of classical DNA damage response pathways and cell-cycle arrest [12,13]. Metabolic stress, including chronic hyperglycaemia and altered glucose utilisation, is also a common feature of ageing and metabolic disease, profoundly shaping microglial inflammatory tone and bioenergetic state [14,15]. Despite their coexistence in ageing, genotoxic and metabolic stress engage distinct upstream signalling pathways, raising the possibility that they drive distinct senescence programmes with divergent cellular and inflammatory outcomes.

Importantly, senescence is increasingly recognised as a spectrum of stable cellular states rather than a singular endpoint. The initiating stressor strongly influences downstream senescence profiles, including differential engagement of cell-cycle arrest mechanisms, mitochondrial adaptation, chromatin organisation and SASP composition [16–18]. Genotoxic stress typically activates a rapid p53-dependent response associated with robust DNA damage signalling, whereas metabolic and nucleolar stress have been linked to p16 induction and alternative senescence trajectories that may stabilise long-term inflammatory phenotypes [19,20]. However, how these mechanistically distinct stressors shape senescence-associated remodelling in human microglia remains poorly understood.

Mitochondrial dysfunction represents another central hallmark of microglial ageing and senescence. Senescent cells frequently exhibit mitochondrial dysfunction, impaired mitochondrial quality control, altered mitochondrial turnover, disrupted redox homeostasis and resistance to apoptotic execution despite evidence of apoptotic priming [21]. In microglia, dysfunctional mitochondria can reinforce inflammatory signalling and sustain senescence-associated phenotypes, thereby perpetuating neuroinflammation. The persistence of senescent, apoptosis-resistant microglia is therefore likely to represent a detrimental and progressive feature of brain ageing.

Despite growing evidence linking neuroinflammation to microglial senescence and ageing, the underlying mechanisms remain incompletely defined, particularly in human systems. Studies of primary human microglia are limited by tissue availability and culture constraints and much existing knowledge is derived from rodent models that do not fully capture human microglial biology [22,23]. Immortalised human microglial cell lines, such as HMC3, provide a convenient and reproducible platform for mechanistic studies. Although immortalisation and the absence of the complete central nervous system environment limit physiological fidelity, HMC3 cells nonetheless provide a robust model: they express key microglial markers, respond to inflammatory stimuli and enable experimentally controlled analysis of stress-induced senescence pathways relevant to human microglia [24,25].

In this study, we investigated how chronic genotoxic and metabolic stress differentially induce senescence-associated phenotypes in human microglia using the HMC3 cell model. Using doxorubicin-induced (Dox) genotoxic stress and chronic high-glucose (HG) exposure as complementary models of ageing-associated stress, we examined morphological remodelling, activation of canonical senescence pathways, mitochondrial and apoptotic responses and the emergence of stress-specific inflammatory signatures. By directly comparing these distinct stressors, this study provides insight into how heterogeneous senescence programmes in microglia may differentially contribute to chronic neuroinflammation and age-related neurodegenerative diseases.

## Methods

### Cell culture

The study was performed using the commercially available human microglial cell line HMC3 (AddexBio, USA; #C0005019). HMC3 cells were cultured in Eagle’s Minimum Essential Medium (EMEM) supplemented with 10% fetal bovine serum (FBS) and 1% sodium pyruvate solution (hereafter referred to as complete medium) at 37 °C in a humidified atmosphere containing 5% CO₂. When cells reached 70–80% confluence, they were trypsinised using trypsin–EDTA for 5 min and collected in fresh complete medium. The cell suspension was centrifuged at 100 × g for 5 min at room temperature (RT), the supernatant was discarded, and the cell pellet was resuspended in complete medium for subsequent experiments.

### Induction of senescence

HMC3 cells (passages 15–22) were seeded at a density of 1 × 10⁴ cells/cm² unless otherwise stated and stress stimuli were applied after 24 h.

For induction of genotoxic stress, doxorubicin (Dox) (Cell Signaling Technology, #5927) was dissolved in water and diluted in complete medium to a final concentration of 50 nM. Cells were treated for 48 h, washed twice with Dulbecco’s phosphate-buffered saline (DPBS; Thermo Fisher Scientific), and cultured for an additional 24 h in fresh complete medium.

For induction of metabolic stress, cells were cultured in complete medium supplemented with 200 mM D-glucose (Sigma, #G7021) for 3 days, with complete medium replacement every other day. This concentration was empirically determined to induce robust senescence-associated phenotypes while maintaining cell viability. Lower glucose concentrations resulted in partial responses, including reduced proliferation and modest induction of senescence markers, but did not consistently produce the full phenotype required for downstream analyses (data not shown).

The use of supra-physiological glucose concentrations is consistent with in vitro models of chronic metabolic stress, where elevated levels are required to induce sustained mitochondrial dysfunction and oxidative stress [26,27]. This requirement may also reflect the increased stress tolerance of immortalised microglial cell lines [24,25].

To control for osmotic effects, cells were cultured in complete medium supplemented with 200 mM D-mannitol (Sigma, #M9546) under identical conditions. Untreated cells and cells maintained in medium containing 5.5 mM glucose served as controls. Cells were harvested at the end of the treatment period for subsequent analyses.

### Cell Viability Assay (MTT)

Cell metabolic activity was assessed using the 3-(4,5-dimethylthiazol-2-yl)-2,5-diphenyltetrazolium bromide (MTT) assay (Roche, #11465007001). At the end of treatment, MTT reagent (0.5 mg/mL final concentration) was added to each well, and cells were incubated for 3 h at 37 °C in 5% CO₂. Absorbance was measured at 600 nm using a microplate reader (Omega).

### Senescence associated β-galactosidase (SA-β-gal) activity

Senescence-associated β-galactosidase (SA-β-gal) activity was measured using a senescence β-galactosidase staining kit (Cell Signaling, #9860) according to the manufacturer’s instructions. Briefly, cells were seeded at a density of 1 × 10⁵ cells per well in 6-well plates. Following treatment, cells were washed once with PBS and fixed for 10 min at room temperature. After washing twice with PBS, cells were incubated with SA-β-gal staining solution (pH 6.0) for 16 h at 37 °C in a CO₂-free incubator. Stained cells were washed with PBS and imaged using a brightfield microscope (EVOS XL Core, Thermo Fisher Scientific). SA-β-gal-positive staining was quantified using CellProfiler (v.4.2.6). Images were subjected to a standardized analysis pipeline in which SA-β-gal-positive staining was identified using automated thresholding and the positive staining area was measured relative to the total image area. Data from individual fields were averaged to generate a single value for each biological replicate prior to statistical analysis.

### Cell morphology staining

Cell morphology was assessed using CellMask™ plasma membrane stain (Invitrogen, #C10046). Following treatment, cells were washed with PBS and incubated with staining solution (1 µL CellMask in 1 mL PBS) for 10 min at 37 °C. Cells were subsequently washed and nuclei counterstained with Hoechst 33342 (Thermo Fisher Scientific, #H3570) for 5 min at 37 °C. After washing, cells were imaged immediately using a fluorescence microscope (EVOS M5000, Thermo Fisher Scientific). Cell area, perimeter, nuclear area and nuclear perimeter were quantified using CellProfiler (v4.2.6). A minimum of 100 cells per biological replicate were analysed. Measurements from individual cells were averaged to generate a single value for each biological replicate.

### Reverse transcription polymerase chain reaction (RT-qPCR)

Total RNA was isolated using the RNeasy Mini Kit (Qiagen, #74536) according to the manufacturer’s protocol. Complementary DNA (cDNA) was synthesised using the Tetro cDNA Synthesis Kit (Meridian Bioscience, #BIO-65043). Quantitative PCR was performed using GoTaq qPCR Master Mix (Promega, #A6002) on a LightCycler® 480 System (Roche). RPS18 and GAPDH were used as housekeeping genes. Cycle threshold (Ct) values were obtained from triplicate measurements and normalised to the mean Ct of housekeeping genes. Primer sequences are listed in Supplementary Table S1.

### Immunocytochemistry (ICC)

Following treatment, cells were fixed in ice-cold methanol for 10 min at −20 °C and washed three times with ice-cold Hank’s Balanced Salt Solution (HBSS). Cells were permeabilised and blocked in HBSS containing 0.1% Triton X-100 (BioXtra, #T9284), 1% goat serum (Abcam, #AB7481), 2% bovine serum albumin (BSA, Sigma, #A8022), and 1% milk for 30 min at room temperature. Cells were incubated with primary antibodies overnight at 4 °C, followed by washing and incubation with Alexa Fluor-conjugated secondary antibodies for 1 h at room temperature in the dark. Nuclei were stained with DAPI (1:1000; Invitrogen, #D1306). Cells were mounted using ProLong™ Diamond Antifade Mountant (Invitrogen, #P36970). Images were acquired using an EVOS M5000 microscope (20× and 40× objectives), and representative images were obtained using a Zeiss confocal microscope. Image analysis was performed using CellProfiler v4.2.6. A custom CellProfiler pipeline was used for automated image analysis. Nuclei were segmented using DAPI staining and automated thresholding. Mean fluorescence intensity was quantified within the nuclear compartment for nuclear markers and within the cytoplasmic compartment for cytoplasmic markers. Cell-level measurements were averaged per image, and image-level values were subsequently averaged to generate a single value for each biological replicate for statistical analysis. Antibody details are provided in Supplementary Table S2.

### MitoTracker^TM^ Green FM staining

Mitochondrial mass was assessed using MitoTracker™ Green FM (Thermo Fisher Scientific, #M7514). A 1 mM stock solution was prepared in anhydrous DMSO and stored at −20 °C.

For staining, the dye was diluted to a final concentration of 100 nM in pre-warmed complete medium. Following treatment, cells were incubated with the staining solution for 30 min at 37 °C in 5% CO₂. Cells were washed and nuclei counterstained with Hoechst 33342 (Thermo Fisher Scientific, #H3570) prior to imaging. Images were acquired using an EVOS M5000 microscope with identical acquisition settings across all conditions. Mitochondrial mass was quantified using CellProfiler, based on mitochondrial area and fluorescence intensity normalised to control cells.

### Western blot

Cells were lysed in RIPA buffer (50mM Tris-HCl, 150mM NaCl, 0.1% SDS, 1% Triton X-100, 0.5% Sodium Deoxycholate, 1mM EDTA, 100mM NaF, 1mM NaOV; pH 7.4) supplemented with protease inhibitors. Protein concentration was determined using the BCA assay (Thermo Fisher Scientific, #23227). Equal amounts of protein (10 µg per sample) were separated by SDS-PAGE (NuPAGE 4–12% Bis-Tris gels) and transferred onto nitrocellulose membranes. Membranes were blocked in 5% skimmed milk in TBS-T (0.05% Tween-20) and incubated with primary antibodies overnight at 4 °C. Following washing, membranes were incubated with HRP-conjugated secondary antibodies for 1 h at room temperature. Protein bands were detected using chemiluminescent substrate (Millipore, #WBKL S0100) and imaged using an iBright FL1000 system. Densitometric analysis was performed using Fiji (v1.53), and protein levels were normalised to total protein staining (Ponceau S, Sigma, #P3504). Antibody details are provided in Supplementary Table S3.

### Statistics analysis

Statistical analyses were performed using GraphPad Prism v11.0.1. Data are presented as mean ± SEM. Data normality was assessed using the Shapiro–Wilk. Statistical differences between two groups were analysed using an unpaired two-tailed Student’s t-test. For comparisons involving more than two groups, one-way analysis of variance (ANOVA) followed by Tukey’s multiple comparisons test was used. Differences were considered statistically significant at p < 0.05. All experiments were performed with five independent biological replicates, except the cell viability (MTT) experiments, which were performed with three independent biological replicates. Technical replicates were averaged prior to statistical analysis. Figures were prepared using Inkscape.

## Results

### Chronic genotoxic and metabolic stress induce senescence-associated morphological remodelling in human microglia cells

To determine whether chronic genotoxic or metabolic stress induces senescence-associated morphological alterations in human microglia, HMC3 cells were exposed to Dox and HG using two independent experimental timelines (**Fig. 1a-b**). These paradigms were selected to model stressors relevant to ageing-associated microglial dysfunction. To control for potential osmotic effects, cells treated with 200 mM HG were compared with those exposed to an equimolar concentration of mannitol (Mtl). Control cells were maintained under standard culture conditions and processed in parallel.

**Fig. 1.**
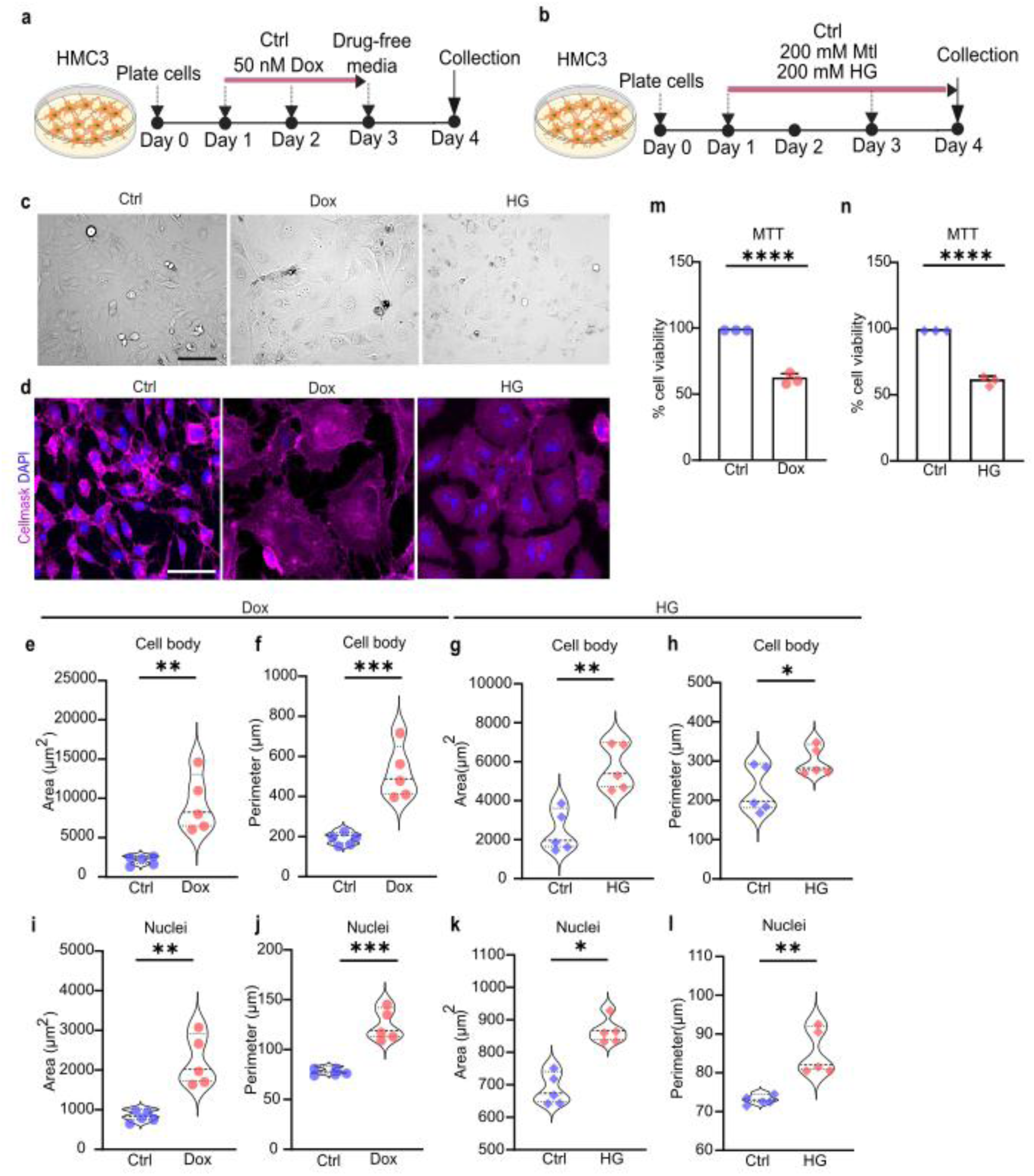
Chronic genotoxic and metabolic stress induce senescence-associated morphological remodelling in human microglia cells. **(a)** Schematic representation of the experimental workflow. HMC3 cells were treated with 50 nM doxorubicin (Dox) for 48 hours. Following drug removal on day 3, cells were maintained in drug-free medium for an additional one day to allow senescence-associated phenotypes to develop. Cells were harvested on day 4 for downstream analyses. **(b)** Schematic representation of the experimental workflow. HMC3 cells were treated with 200 mM high glucose (HG) and equimolar mannitol (Mtl) for 72 hours with a full medium change every other day. Cells were harvested on day 4 for downstream analyses. **(c)** Representative brightfield images showing microglial morphology of Ctrl and Dox or HG-treated HMC3 cells. Scale bar: 100 µm **(d)** Representative images of morphology staining showing microglial morphology of Ctrl and Dox or HG-treated HMC3 cells. Scale bar: 100 µm **(e-h)** Quantification of cell body areas and perimeters in Ctrl and Dox or HG-treated HMC3 cells. Values represent mean + SEM from *n* = 5 biological replicates, \**p* < 0.05; \*\**p* < 0.01 \*\*\**p* < 0.001, unpaired two-tailed Student’s *t* test. **(i-l)** Quantification of nuclear areas and perimeters in Ctrl and Dox or HG-treated HMC3 cells. Values represent mean + SEM from *n* = 5 biological replicates, *p* < 0.05; \*\**p* < 0.01 \*\*\**p* < 0.001, unpaired two-tailed Student’s *t* test. **(m-n)** MTT assay showing the effects of Dox and HG treatment on HMC3 cell metabolic activity. Values represent mean + SEM from *n* = 3 biological replicates, \*\*\*\**p* < 0.0001, unpaired two-tailed Student’s *t* test.

Brightfield imaging revealed marked morphological remodelling in HMC3 cells following exposure to both Dox and HG relative to control cultures (**Fig. 1c**). Control cells exhibited a small, compact soma with well-defined cellular boundaries, consistent with a resting microglial phenotype. In contrast, both senescence-inducing paradigms resulted in enlarged and flattened cells with irregular boundaries. Dox-treated cells displayed pronounced cellular hypertrophy and loss of clearly defined edges, whereas HG-treated cells exhibited increased cell size and flattening, with greater heterogeneity across the population. Notably, HG-treated cells also showed a more vacuolated and occasionally rounded morphology, suggestive of altered metabolic state. In contrast, cells exposed to equimolar Mtl displayed only minor deviations from control cultures, with slight changes in cell size, and remained clearly distinct from the pronounced alterations observed under HG conditions, indicating that these effects were not driven by osmotic stress (**Fig. S1a**).

To further characterise alterations in microglial structural organisation, morphology-specific staining was performed (**Fig. 1d**). Control cells displayed uniform cytoplasmic organisation and clearly delineated cellular boundaries. Cells exposed to Mtl exhibited a comparable morphology, closely resembling control cultures, with only a slight increase in cell size while preserving overall cellular architecture (**Fig. S2b**). In contrast, Dox– and HG-treated cells demonstrated disrupted cellular organisation, including extensive cytoplasmic spreading and irregular borders. These features are consistent with a senescence-associated morphological phenotype and corroborate observations from brightfield imaging.

Quantitative morphometric analysis confirmed significant enlargement of the microglial cell body following senescence induction. Both Dox and HG treatments resulted in a significant increase in cell body area (4.4-fold, 2.3-fold, respectively) compared with controls **(Fig. 1e, g**), accompanied by a corresponding increase in cell body perimeter (2.7-fold, 1.3-fold, respectively) (**Fig. 1f, h**). In contrast, Mtl-treated cells also showed a modest but significant increase in cell body area relative to controls, whereas perimeter remained unchanged, indicating that the morphometric alterations observed under HG conditions cannot be attributed solely to osmotic stress (**Fig. S2c–d**). Direct comparison between the two stress paradigms revealed that Dox exposure induced a significantly greater increase in both cell body area and perimeter than HG treatment, suggesting a more robust morphological response to genotoxic stress (**Fig. S3a–b**).

Given that nuclear hypertrophy is a well-established feature of cellular senescence, nuclear morphometric parameters were also assessed. Both Dox– and HG-treated HMC3 cells exhibited a significant increase in nuclear area (2.6-fold, 1.25-fold respectively) relative to controls (**Fig. 1i, k**), accompanied by a corresponding increase in nuclear perimeter (1.6-fold, 1.2-fold respectively) (**Fig. 1j, l**). In contrast, nuclei from Mtl-treated cells did not differ significantly from controls in either area or perimeter, consistent with the absence of osmotic stress-induced senescent changes (**Fig. S2e–f**). As observed for whole-cell metrics, Dox treatment induced a greater magnitude of nuclear enlargement than HG exposure, indicating differential effects of these stressors on nuclear architecture (**Fig. S3c–d**).

Following morphological characterisation, metabolic activity was assessed to determine the impact of chronic stress on microglial function. MTT analysis demonstrated a significant reduction in mitochondrial metabolic activity following both Dox and HG treatment relative to control conditions. Dox exposure reduced metabolic activity to 59.2% of control levels, whereas HG treatment resulted in a comparable reduction to 62.0% (**Fig. 1m, n**). In contrast, Mtl-treated cells retained significantly higher metabolic activity, at approximately 78.6% relative to controls (**Fig. S2a**). This partial reduction, compared with the more pronounced decrease observed under HG conditions, indicates that the effect of HG is not solely attributable to osmotic stress.

These comparable reductions indicate that both treatments impose sustained cellular stress sufficient to impair mitochondrial metabolic function without inducing complete loss of cell viability, thereby enabling downstream analysis of senescence-associated phenotypes.

### Distinct engagement of p53-p21 and p16 pathways in stress-induced microglial senescence

To further confirm acquisition of a senescent phenotype in response to chronic stress, SA-β-gal activity was assessed in HMC3 cells following exposure to Dox and HG (**Fig. 2a, c**). Both treatments resulted in a significant increase in the proportion of SA-β-gal-positive area compared with control, providing functional evidence of senescence induction (**Fig. 2b, d**). The magnitude of SA-β-gal activity was comparable between Dox– and HG-treated cells, indicating that both genotoxic and metabolic stress are sufficient to induce lysosomal alterations characteristic of senescent microglia (**Fig. S3e**).

**Fig. 2.**
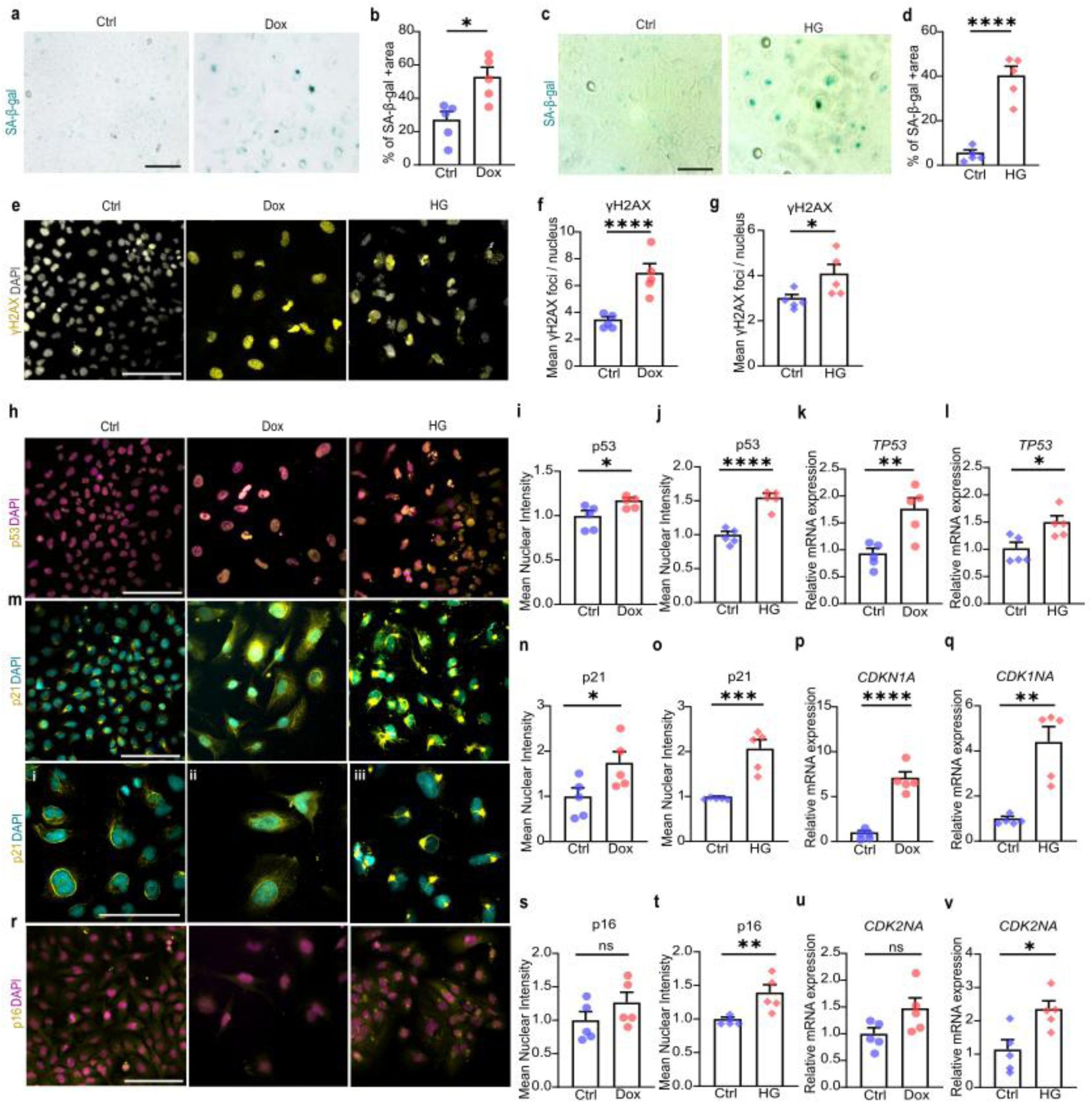
Distinct engagement of p53-p21 and p16 pathways in stress-induced microglial senescence. **(a-d)** Quantification of the area percentage of senescence-associated β-galactosidase staining in Ctrl and Dox or HG-treated HMC3 cells. Values represent mean + SEM from *n* = 5 biological replicates, \**p* < 0.05; \*\*\*\**p* < 0.0001, unpaired two-tailed Student’s *t* test. Scale bar: 100 µm **(e)** Representative images of γH2AX^+^ foci in Ctrl and Dox or HG-treated HMC3 cells. Scale bar: 100 µm **(f-g)** Quantification of the percentage of nuclei with γH2AX+ foci in Ctrl and Dox or HG-treated HMC3 cells. Values represent mean + SEM from *n* = 5 biological replicates, \**p* < 0.05; \*\*\*\**p* < 0.0001, unpaired two-tailed Student’s *t* test. **(h)** Representative images of p53 immunofluorescence staining in Ctrl and Dox or HG-treated HMC3 cells. Scale bar: 100 µm **(i-j)** Quantification of mean nuclear p53 fluorescence intensity in Ctrl and Dox or HG-treated HMC3 cells. Values represent mean + SEM from *n* = 5 biological replicates, \**p* < 0.05; \*\*\*\**p* < 0.0001, unpaired two-tailed Student’s *t* test. **(k-l)** Relative mRNA expression levels of *TP53* in Ctrl and Dox or HG-treated HMC3 cells measured by RT–qPCR. Values represent mean + SEM from n = 5 biological replicates, normalised to the housekeeping genes *RPS18* and *GAPDH*, \**p* < 0.05; \*\**p* < 0.01, unpaired two-tailed Student’s *t* test. **(m)** Representative images of p21 immunofluorescence staining in Ctrl and Dox or HG-treated HMC3 cells. Scale bar: 100 µm **(n-o)** Quantification of mean nuclear p21 fluorescence intensity in Ctrl and Dox or HG-treated HMC3 cells. Values represent mean + SEM from *n* = 5 biological replicates, \**p* < 0.05; \*\*\**p* < 0.001, unpaired two-tailed Student’s *t* test. **(p-q)** Relative mRNA expression levels of *CDKN1A* in Ctrl and Dox or HG-treated HMC3 cells measured by RT–qPCR. Values represent mean + SEM from n = 5 biological replicates, normalised to the housekeeping genes *RPS18* and *GAPDH*, \*\**p* < 0.01; \*\*\*\**p* < 0.0001, unpaired two-tailed Student’s *t* test. **(r)** Representative images of p16 immunofluorescence staining in Ctrl and Dox or HG-treated HMC3 cells. Scale bar: 100 µm **(s-t)** Quantification of mean nuclear p16 fluorescence intensity in Ctrl and Dox or HG-treated HMC3 cells. Values represent mean + SEM from *n* = 5 biological replicates, ns; not significant, \*\**p* < 0.01, unpaired two-tailed Student’s *t* test. **(u-v)** Relative mRNA expression levels of *CDKN2A* in Ctrl and Dox or HG-treated HMC3 cells measured by RT–qPCR. Values represent mean + SEM from n = 5 biological replicates, normalised to the housekeeping genes *RPS18* and *GAPDH*, ns; not significant, \**p* < 0.05, unpaired two-tailed Student’s *t* test.

Given that persistent DNA damage is a key driver of cellular senescence, DNA damage responses were next evaluated by quantifying γH2AX foci formation (**Fig. 2e**). Both Dox– and HG-treated HMC3 cells exhibited a significant increase in nuclear γH2AX foci relative to control cells, indicating accumulation of DNA damage under both chronic stress conditions (**Fig. 2f, g**). Notably, γH2AX induction was significantly greater following Dox exposure than HG treatment, consistent with the direct genotoxic effects of Dox (**Fig. S3f**). These findings indicate that, although both stress paradigms activate DNA damage-associated signalling, genotoxic stress elicits a more robust DNA damage response than metabolic stress.

To determine whether these senescence-associated changes were accompanied by activation of canonical cell cycle arrest pathways, expression of p53, p21 and p16 was examined at both transcript and protein levels using quantitative PCR (qPCR) and immunocytochemistry (ICC). qPCR analysis revealed significant upregulation of *TP53* (p53) and *CDKN1A* (p21) mRNA in HMC3 cells following both Dox and HG treatment compared with controls (**Fig. 2k–l, p–q**). Notably, cells exposed to equimolar Mtl also exhibited a significant increase in p53 and p21 transcript levels relative to controls, indicating that activation of these cell cycle arrest pathways may, at least in part, be influenced by osmotic stress rather than being exclusively driven by metabolic perturbation under HG conditions (**Fig. S2g–h**). In agreement with these transcriptional findings, ICC demonstrated increased p53 and p21 protein immunoreactivity in both Dox– and HG-treated cells (**Fig. 2h–j, m–o**).

Notably, p21 displayed distinct subnuclear localisation patterns depending on the nature of the stressor. Qualitative assessment indicated predominantly pan-nuclear (nucleoplasmic) localisation in Dox-treated cells, consistent with activation of a canonical DNA damage-induced cell cycle checkpoint (**Fig. 2m (ii)**). In contrast, HG treatment induced pronounced peripheral nuclear accumulation of p21 compared with control (**Fig. 2m (i, iii)**). This differential subnuclear distribution suggests engagement of stimulus-specific cell cycle arrest mechanisms, with HG preferentially activating metabolic stress-associated pathways linked to senescence, whereas Dox predominantly induces a classical genotoxic stress-driven p53-p21 response.

This coordinated induction of p53 and p21 at both transcript and protein levels across treatment conditions indicates activation of a canonical p53-p21 signalling axis consistent with stress-induced cell cycle arrest. Despite the distinct nature of the applied stressors, both paradigms converged on comparable p53 and p21 upregulation, suggesting shared upstream regulatory mechanisms.

In contrast to p53 and p21, analysis of p16 expression revealed a stressor-dependent response. Under HG conditions, p16 expression was significantly increased at both mRNA and protein levels compared with control cells, consistent with engagement of a p16-associated senescence programme (**Fig. 2r, t, v**). By comparison, Dox treatment induced modest and variable changes in p16 expression that did not reach statistical significance (**Fig. 2r, s, u**). Notably, qPCR analysis showed that equimolar Mtl exposure induced *CDKN2A* (p16) mRNA expression relative to controls, suggesting that osmotic stress may contribute to HG-mediated p16 activation (**Fig. S2i**).

Collectively, these findings demonstrate that chronic genotoxic and metabolic stress promote senescence-like phenotypes in human microglia, characterised by increased SA-β-gal activity, persistent DNA damage signalling, and activation of growth arrest pathways. While both stress paradigms converge on p53–p21 dependent cell cycle arrest, metabolic stress is associated with enhanced p16 expression and a distinct peripheral nuclear p21 localisation pattern. These differences suggest that, rather than representing variations in stress severity, genotoxic and metabolic stress drive fundamentally distinct senescence architectures, reflecting divergence in the underlying regulatory and functional programmes.

### Dox-induced stress induces apoptotic priming and mitochondrial remodelling without execution-phase apoptosis

To assess whether Dox-induced stress engages apoptotic pathways in microglia, we examined key regulators of intrinsic apoptosis at both transcriptional and protein levels. Quantitative PCR analysis revealed no significant changes in mRNA expression of *CASP3* or the anti-apoptotic regulator *BCL2* following Dox exposure (**Fig. 3a-b**). In contrast, western blot analysis demonstrated a significant increase in procaspase-3 protein levels compared with control conditions (**Fig. 3c, d**). Despite this increase, cytochrome c protein levels remained unchanged, indicating that mitochondrial outer membrane permeabilisation and downstream caspase activation were not initiated (**Fig. 3c, e**).

**Fig. 3.**
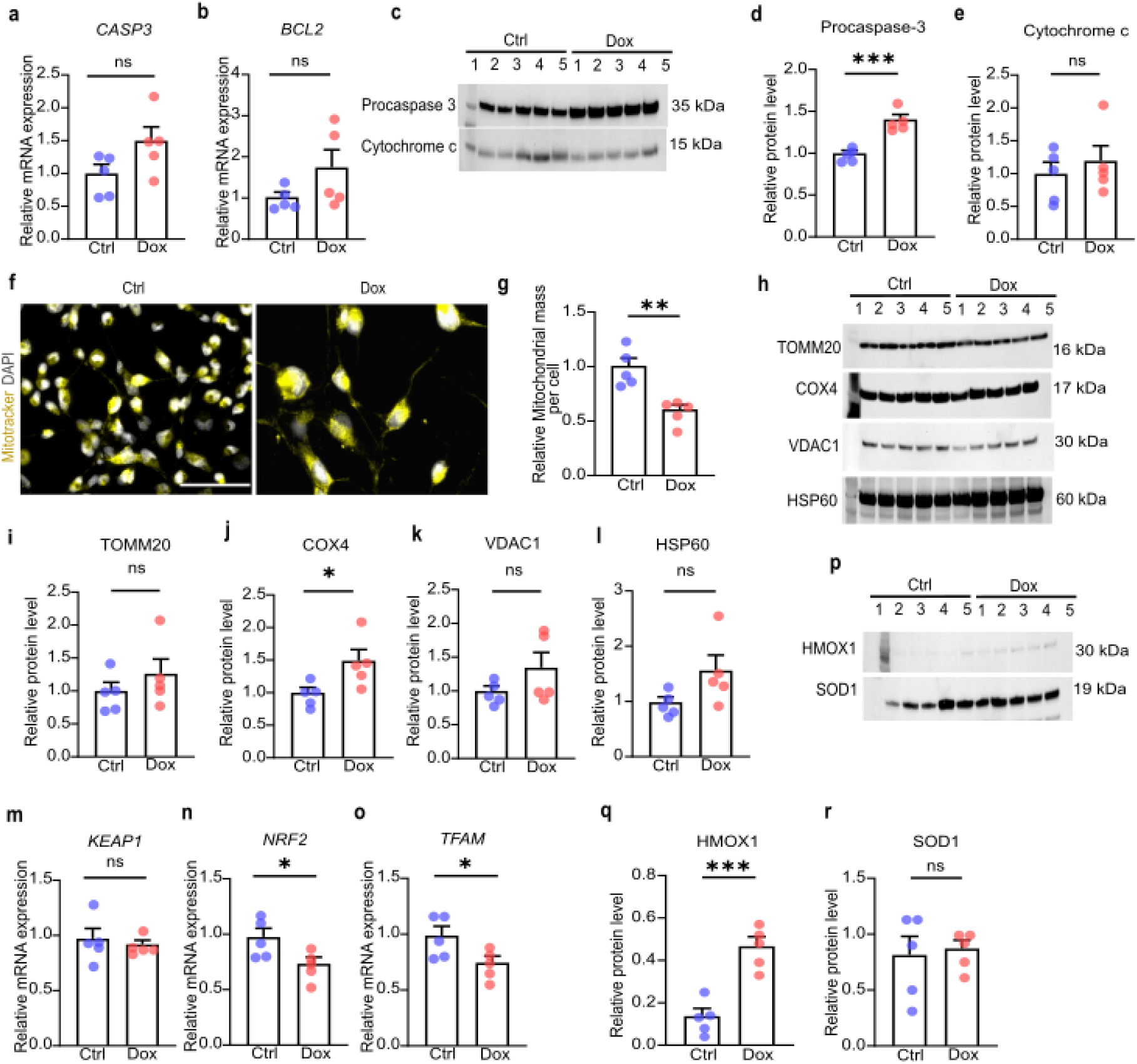
Dox-induced stress induces apoptotic priming and mitochondrial remodelling without execution-phase apoptosis. **(a)** Relative mRNA expression levels of *CASP3* in Ctrl and Dox-treated HMC3 cells measured by RT–qPCR. Values represent mean + SEM from n = 5 biological replicates, normalised to the housekeeping genes *RPS18* and *GAPDH*, ns; not significant, unpaired two-tailed Student’s *t* test. **(b)** Relative mRNA expression levels of *BCL2* in Ctrl and Dox-treated HMC3 cells measured by RT–qPCR. Values represent mean + SEM from n = 5 biological replicates, normalised to the housekeeping genes *RPS18* and *GAPDH*, ns; not significant, unpaired two-tailed Student’s *t* test. **(c)** Representative western blots showing the expression of apoptosis related proteins procaspase-3 and cytochrome c in Ctrl and Dox-treated HMC3 cells. Shown blots are representative of n=5 biological replicates. **(d-e)** Quantification of procaspase-3 and cytochrome c protein levels in Ctrl and Dox-treated HMC3 cells, normalised to total protein. Values represent mean + SEM from *n* = 5 biological replicates, ns; not significant, \*\*\**p* < 0.001, unpaired two-tailed Student’s *t* test. **(f)** Representative images of Mitotracker staining in Ctrl and Dox-treated HMC3 cells. Scale bar: 100 µm **(g)** Quantification of the relative mitochondrial mass per cell in Ctrl and Dox-treated HMC3 cells. Values represent mean + SEM from *n* = 5 biological replicates, \*\**p* < 0.01, unpaired two-tailed Student’s *t* test. **(h)** Representative western blots showing the expression of mitochondrial proteins TOMM20, COX4, VDAC1 and HSP60 in Ctrl and Dox-treated HMC3 cells. Shown blots are representative of n=5 biological replicates. **(i-l)** Quantification of TOMM20, COX4, VDAC1, HSP60 protein levels in Ctrl and Dox-treated HMC3 cells, normalised to total protein. Values represent mean + SEM from *n* = 5 biological replicates, ns; not significant, \**p* < 0.05, unpaired two-tailed Student’s *t* test. **(m-o)** Relative mRNA expression levels of *KEAP1, NRF2, TFAM* in Ctrl and Dox-treated HMC3 cells measured by RT–qPCR. Values represent mean + SEM from n = 5 biological replicates, normalised to the housekeeping genes *RPS18* and *GAPDH*, \**p* < 0.05, ns; not significant, unpaired two-tailed Student’s *t* test. **(p)** Representative western blots showing the expression of antioxidant response proteins HMOX1 and SOD1 in Ctrl and Dox-treated HMC3 cells. Shown blots are representative of n=5 biological replicates. **(q-r)** Quantification of HMOX1 and SOD1 protein levels in Ctrl and Dox-treated HMC3 cells, normalised to total protein. Values represent mean + SEM from *n* = 5 biological replicates, ns; not significant, \*\*\*\**p* < 0.0001, unpaired two-tailed Student’s *t* test.

To further characterise mitochondrial responses to Dox-induced stress, mitochondrial mass was first assessed using MitoTracker staining (**Fig. 3f**). Quantitative analysis revealed a significant reduction in mitochondrial mass in Dox-treated microglia (**Fig. 3g**). Consistent with these findings, western blot analysis of mitochondrial proteins showed no significant changes in the abundance of the outer mitochondrial membrane protein VDAC1 or the mitochondrial import receptor TOMM20 (**Fig. 3h, i, k**). Similarly, expression of the mitochondrial matrix chaperone HSP60 remained unchanged (**Fig. 3h, l**). Despite the reduction in mitochondrial mass, Dox-treated cells exhibited selective upregulation of cytochrome c oxidase subunit IV (COX4), a core component of the electron transport chain, while other mitochondrial markers remained unchanged. This suggests differential regulation of mitochondrial components rather than uniform mitochondrial loss (**Fig. 3h, j**).

To further investigate mitochondrial stress responses, we examined key regulators of antioxidant signalling and mitochondrial biogenesis, including KEAP1, a negative regulator of NRF2, NRF2 itself, a master transcriptional regulator of cellular antioxidant responses and TFAM, which governs mitochondrial transcription and biogenesis. Despite the reduction in mitochondrial mass, transcriptional analysis revealed no significant change in *KEAP1* expression, accompanied by significant downregulation of *NRF2* and *TFAM* in Dox-treated cells compared with controls (**Fig. 3m–o**). These findings suggest impaired activation of stress-responsive antioxidant pathways and reduced mitochondrial biogenesis signalling. Consistent with this, the transcriptional profile does not support restoration of mitochondrial content, indicating a failure of compensatory mitochondrial recovery under chronic genotoxic stress.

In line with these observations, protein analysis revealed that the NRF2 target HMOX1 was significantly upregulated in Dox-treated microglia, whereas SOD1 levels remained unchanged (**Fig. 3p–r**). This pattern suggests selective activation of NRF2 target genes, potentially reflecting a context-dependent or partial antioxidant response rather than a fully coordinated transcriptional programme.

Together, these data indicate that Dox-induced genotoxic stress results in a reduction in mitochondrial mass and establishes an apoptosis-primed state in microglia, as evidenced by increased procaspase-3 levels, without progression to mitochondrial apoptotic execution. In contrast to a compensatory adaptive response, *NRF2* and *TFAM* expression were significantly downregulated, indicating impaired antioxidant and mitochondrial regulatory signalling. Consistent with this, the transcriptional profile does not support restoration of mitochondrial mass or global antioxidant capacity. The selective upregulation of HMOX1, alongside unchanged SOD1 levels, further supports a partial and context-dependent stress response rather than a fully coordinated transcriptional programme. This phenotype is consistent with mitochondrial dysfunction coupled to defective compensatory adaptation, a feature of stress-induced senescence that may promote persistence of a pro-inflammatory microglial state rather than cell elimination.

### High glucose induces mitochondrial remodelling and transcriptional stress responses without apoptotic execution, independent of osmotic stress

In contrast to genotoxic stress, HG-induced stress showed a prominent transcriptional response without activation of apoptotic execution pathways. Quantitative PCR analysis revealed significant upregulation of *CASP3* and the anti-apoptotic regulator *BCL2* mRNA following HG exposure, indicating engagement of stress-responsive, apoptosis-related gene programmes (**Fig. 4a, b**). Under osmotic control conditions, equimolar Mtl exposure also induced a significant increase in CASP3 expression; however, BCL2 expression remained unaltered, suggesting that osmotic stress alone is insufficient to induce the coordinated pro-survival transcriptional response observed under HG conditions (**Fig. S2j–k**). In addition, western blot analysis demonstrated no significant changes in procaspase-3 or cytochrome c protein levels in HG-treated cells compared with controls, indicating that transcriptional induction does not translate into activation of the mitochondrial apoptotic cascade (**Fig. 4c– e**).

**Fig. 4.**
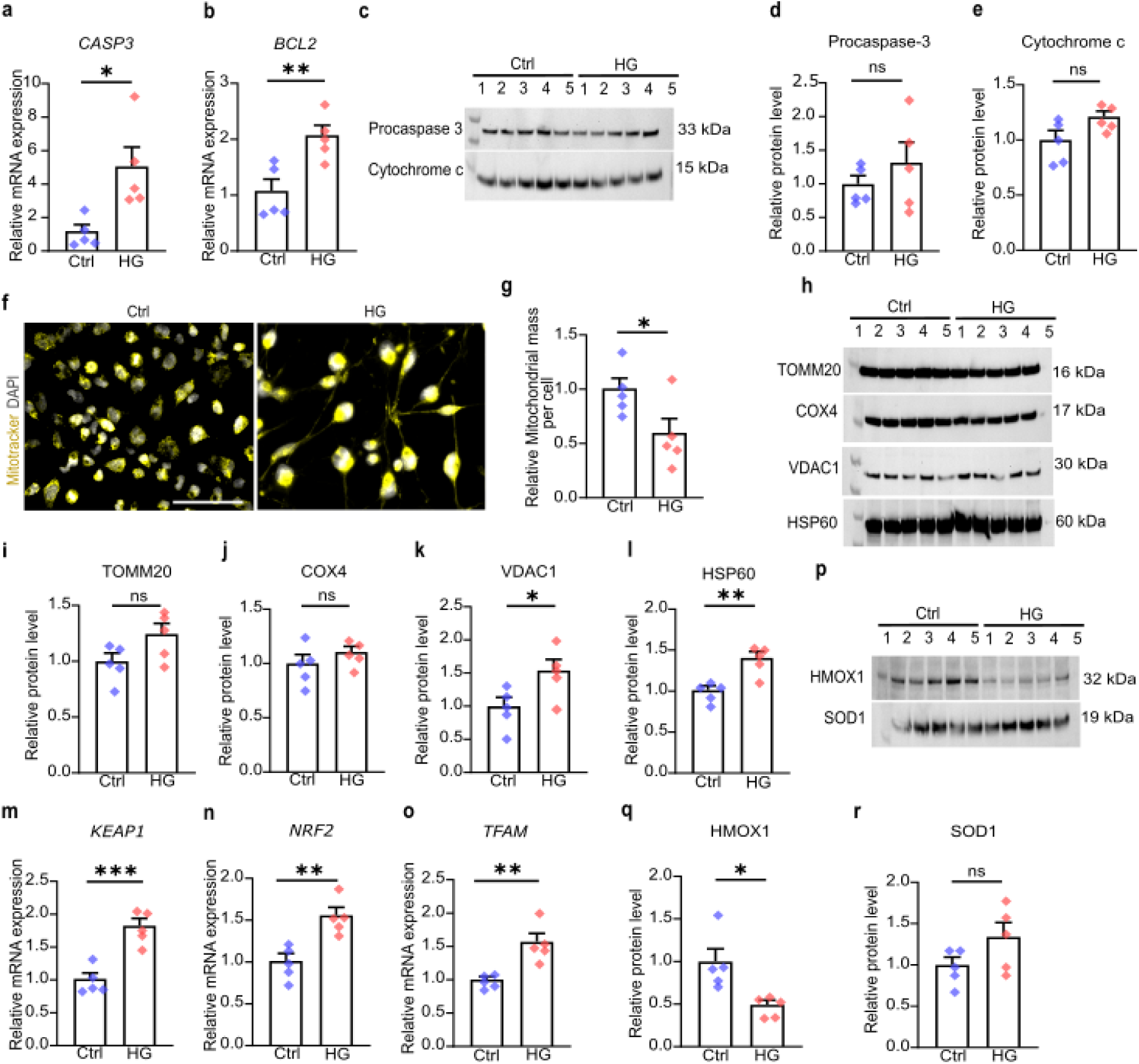
High glucose induces mitochondrial remodelling and transcriptional stress responses without apoptotic execution, independent of osmotic stress. **(a)** Relative mRNA expression levels of *CASP3* in Ctrl and HG-treated HMC3 cells measured by RT–qPCR. Values represent mean + SEM from n = 5 biological replicates, normalised to the housekeeping genes *RPS18* and *GAPDH*, \**p* < 0.05, unpaired two-tailed Student’s *t* test. **(b)** Relative mRNA expression levels of *BCL2* in Ctrl and HG-treated HMC3 cells measured by RT–qPCR. Values represent mean + SEM from n = 5 biological replicates, normalised to the housekeeping genes *RPS18* and *GAPDH*, \*\**p* < 0.01, unpaired two-tailed Student’s *t* test. **(c)** Representative western blots showing the expression of apoptosis related proteins procaspase-3 and cytochrome c in Ctrl and HG-treated HMC3 cells. Shown blots are representative of n=5 biological replicates. **(d-e)** Quantification of procaspase-3 and cytochrome c protein levels in Ctrl and HG-treated HMC3 cells, normalised to total protein. Values represent mean + SEM from *n* = 5 biological replicates, ns; not significant, unpaired two-tailed Student’s *t* test. **(f)** Representative images of Mitotracker staining in Ctrl and HG-treated HMC3 cells. Scale bar: 100 µm **(g)** Quantification of the relative mitochondrial mass per cell in Ctrl and HG-treated HMC3 cells. Values represent mean + SEM from *n* = 5 biological replicates, \**p* < 0.05, unpaired two-tailed Student’s *t* test. **(h)** Representative western blots showing the expression of mitochondrial proteins TOMM20, COX4, VDAC1 and HSP60 in Ctrl and HG-treated HMC3 cells. Shown blots are representative of n=5 biological replicates. **(i-l)** Quantification of TOMM20, COX4, VDAC1, HSP60 protein levels in Ctrl and HG-treated HMC3 cells, normalised to total protein. Values represent mean + SEM from *n* = 5 biological replicates, ns; not significant, \**p* < 0.05; \*\**p* < 0.01, unpaired two-tailed Student’s *t* test. **(m-o)** Relative mRNA expression levels of *KEAP1, NRF2, TFAM* in Ctrl and HG-treated HMC3 cells measured by RT–qPCR. Values represent mean + SEM from n = 5 biological replicates, normalised to the housekeeping genes *RPS18* and *GAPDH*, \*\**p* < 0.01; \*\*\**p* < 0.001, unpaired two-tailed Student’s *t* test. **(p)** Representative western blots showing the expression of antioxidant response proteins HMOX1 and SOD1 in Ctrl and HG-treated HMC3 cells. Shown blots are representative of n=5 biological replicates. **(q-r)** Quantification of HMOX1 and SOD1 protein levels in Ctrl and HG-treated HMC3 cells, normalised to total protein. Values represent mean + SEM from *n* = 5 biological replicates, ns; not significant, \**p* < 0.05, unpaired two-tailed Student’s *t* test.

Analysis of mitochondrial markers revealed a distinct pattern of mitochondrial remodelling under HG conditions. MitoTracker staining and quantitative analysis demonstrated a significant reduction in mitochondrial mass in HG-treated microglia (**Fig. 4f, g**). In contrast, western blot analysis showed significant upregulation of VDAC1 protein, whereas levels of TOMM20 and COX4 remained unchanged (**Fig. 4h–k**). Notably, HSP60 protein expression was significantly increased, indicative of enhanced mitochondrial proteotoxic stress (**Fig. 4h, l**).

Unlike Dox-induced stress, the reduction in mitochondrial mass under HG conditions was accompanied by coordinated transcriptional activation of redox and mitochondrial regulatory pathways. Quantitative PCR analysis demonstrated significant upregulation of *KEAP1, NRF2* and *TFAM*, suggesting engagement of oxidative stress-responsive signalling and TFAM-associated mitochondrial transcriptional programmes (**Fig. 4m–o**). In contrast, Mtl-treated cells did not exhibit significant changes in *NRF2* expression, indicating that activation of redox-sensitive transcriptional pathways is specific to metabolic stress rather than osmotic imbalance (**Fig. S2l**).

However, western blot analysis revealed a significant reduction in the NRF2 target protein HMOX1 in HG-treated microglia, while SOD1 expression showed only a modest, non-significant increase (**Fig. 4p–r**). This divergence suggests a dysregulated or incomplete antioxidant response, whereby upstream redox sensing and mitochondrial regulatory pathways are transcriptionally activated without corresponding induction of key cytoprotective effectors.

Collectively, these data indicate that HG-induced stress is associated with a reduction in mitochondrial mass alongside activation of redox-sensitive and mitochondrial regulatory transcriptional programmes, in the absence of apoptotic execution. Importantly, the selective upregulation of the anti-apoptotic factor BCL2 under HG, but not Mtl conditions, suggests that metabolic stress promotes an apoptosis-resistant phenotype rather than a purely stress-induced response. This phenotype may involve mitochondrial remodelling processes that favour cell survival, potentially through preservation of mitochondrial integrity and inhibition of cytochrome c release. However, the concomitant reduction in HMOX1 protein expression, despite increased *KEAP1–NRF2–TFAM* transcript levels, indicates an uncoupling between redox-responsive transcriptional activation and downstream antioxidant effector induction.

Taken together (see Table 1), these findings demonstrate that chronic metabolic stress activates upstream redox and mitochondrial regulatory transcriptional responses yet fails to restore mitochondrial content or antioxidant effector expression, consistent with maladaptive mitochondrial remodelling in senescent microglia.

**Table 1.** Differential regulation of mitochondrial and antioxidant pathways under genotoxic and metabolic stress.

| Marker | Dox (Genotoxic Stress) | HG (Metabolic Stress) |
| --- | --- | --- |
| **KEAP1** | No significant change | ↑ Increased |
| **NRF2** | ↓ Downregulated | ↑ Increased |
| **TFAM** | ↓ Downregulated | ↑ Increased |
| **HMOX1** | ↑ Increased | ↓ Decreased |
| **SOD1** | No significant change | No significant change |

### Genotoxic and metabolic stress induce distinct senescence-associated inflammatory profiles in microglia

Cellular senescence has emerged as an important contributor to chronic neuroinflammation through the development of the SASP. To determine whether genotoxic or metabolic stress induces senescence-associated inflammatory signalling in microglia, we examined the expression of key SASP-associated cytokines and chemokines following exposure to Dox and HG.

Heatmap analysis revealed distinct SASP profiles between Dox– and HG-induced senescence (**Fig. 5a–b**). Dox induced a more uniform and robust pro-inflammatory signature, whereas HG led to a more heterogeneous SASP characterised by selective chemokine expression. Network-based representation of SASP components further demonstrated stimulus-specific organisation of inflammatory pathways in Dox– and HG-treated cells (**Fig. 5c**). Specifically, Dox induced a centralised, cytokine-dominated SASP network, whereas HG promoted a more heterogeneous, multi-pathway response.

**Fig. 5.**
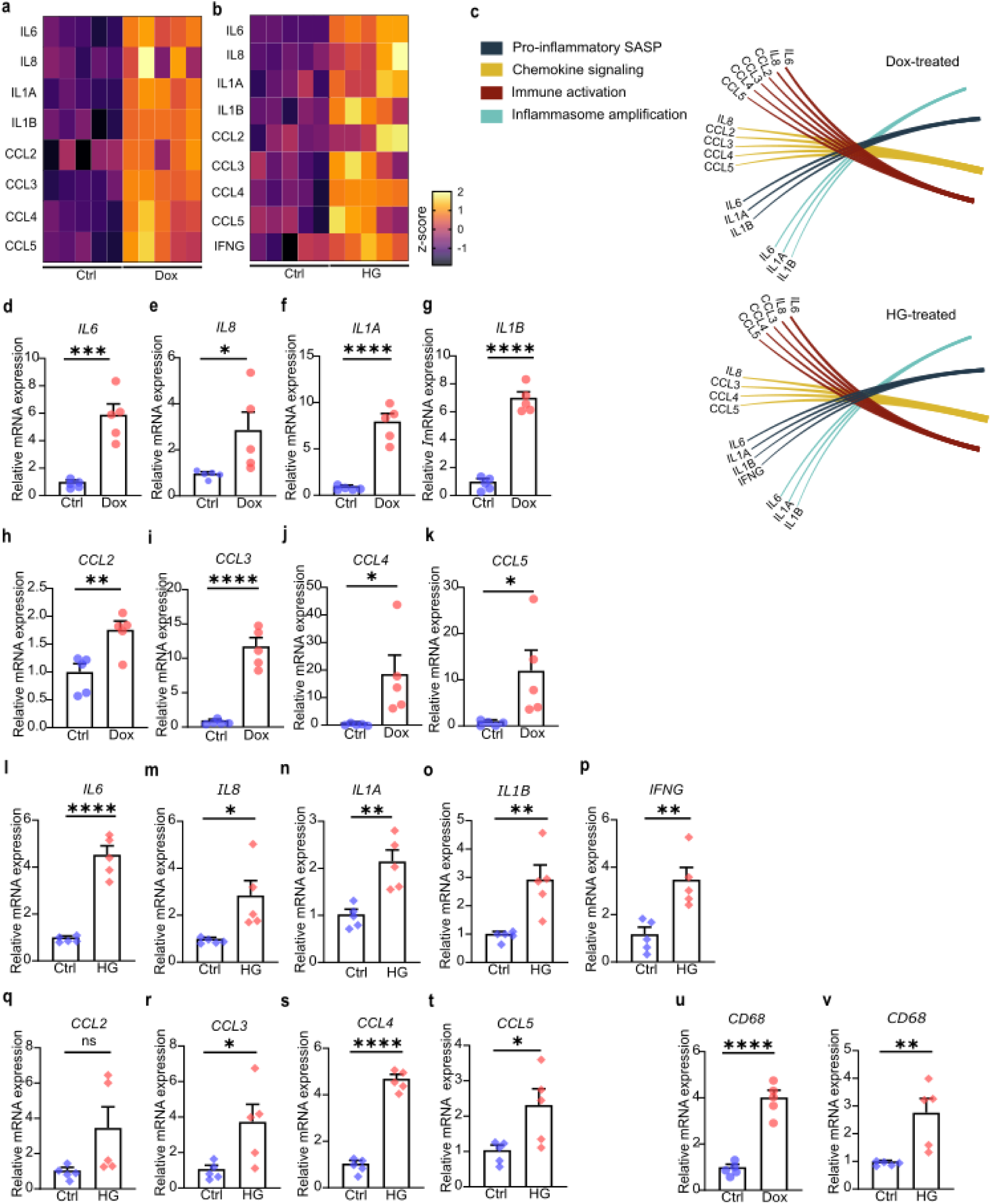
Genotoxic and metabolic stress induce distinct senescence-associated inflammatory profiles in microglia. **(a-b)** Heatmaps showing the expression of SASP in Ctrl and Dox or HG-treated HMC3 cells based on log₂ fold-change (log₂FC) values. Log₂FC values were converted to z-scores for each gene and visualised using GraphPad Prism. Each row represent a gene and each column an individual sample (n=5 Ctrl, Dox or HG-treated) and colours indicate the z-score of each gene across samples. **(c)** Radial plot showing pathway-based functional classification of genes associated with proinflammatory SASP, chemokine signalling, immune activation and inflammasome amplification in Ctrl and Dox or HG-treated HMC3 cells. Genes are grouped according to pathway assignment, with coloured segments indicating each category. **(d-k)** Relative mRNA expression levels of *IL6*, *IL8*, *IL1A*, *IL1B*, *CCL2*, *CCL3*, *CCL4* and *CCL5* in Ctrl and Dox-treated HMC3 cells measured by RT–qPCR. Values represent mean + SEM from n = 5 biological replicates, normalised to the housekeeping genes *RPS18* and *GAPDH*, \**p* < 0.05; \*\**p* < 0.01; \*\*\**p* < 0.001; \*\*\*\**p* < 0.0001, unpaired two-tailed Student’s *t* test. **(l-t)** Relative mRNA expression levels of *IL6*, *IL8*, *IL1A*, *IL1B*, *IFNG*, *CCL2*, *CCL3*, *CCL4* and *CCL5* in Ctrl and HG-treated HMC3 cells measured by RT–qPCR. Values represent mean + SEM from n = 5 biological replicates, normalised to the housekeeping genes *RPS18* and *GAPDH*, ns; not significant, \**p* < 0.05; \*\**p* < 0.01; \*\*\*\**p* < 0.0001, unpaired two-tailed Student’s *t* test. **(u-v)** Relative mRNA expression levels of *CD68*, in Ctrl and Dox or HG-treated HMC3 cells measured by RT–qPCR. Values represent mean + SEM from n = 5 biological replicates, normalised to the housekeeping genes *RPS18* and *GAPDH*, \*\**p* < 0.01; \*\*\*\**p* < 0.0001, unpaired two-tailed Student’s *t* test.

Dox treatment induced a pronounced pro-inflammatory SASP in microglia. Quantitative PCR analysis revealed significant upregulation of canonical SASP cytokines, including *IL6, IL8, IL1B* and *IL1A,* relative to control conditions (**Fig. 5d–g**). In addition, several chemokines involved in immune cell recruitment and amplification of inflammatory signalling, including *CCL2, CCL3, CCL4* and *CCL5*, were significantly upregulated (**Fig. 5h–k**). This broad cytokine-chemokine profile is consistent with a DNA damage-driven senescence response, in which persistent genotoxic stress promotes sustained inflammatory signalling, immune cell recruitment and paracrine reinforcement of senescence rather than apoptotic clearance.

Following SASP induction, Dox-treated microglia also exhibited a marked increase in CD68 expression compared with controls (**Fig. 5u**). Upregulation of CD68 is consistent with a lysosome-enriched, activated microglial phenotype, supporting the presence of sustained inflammatory activation rather than transient stress responses or apoptotic cell loss.

HG exposure likewise resulted in significant upregulation of *IL6, IL8, IL1B* and *IL1A*, indicating activation of a core senescence-associated inflammatory programme in response to metabolic stress (**Fig. 5l–o**). In contrast to the Dox condition, HG treatment led to a significant increase in *IFNG* expression, suggesting preferential engagement of inflammatory pathways linked to metabolic dysregulation and immune signalling (**Fig. 5p**). Analysis of chemokine expression revealed a distinct profile, with no significant change in *CCL2* levels, whereas *CCL3, CCL4* and *CCL5* were significantly upregulated (**Fig. 5q–t**). This pattern indicates activation of selective chemotactic and immunomodulatory signalling pathways under HG conditions, distinct from the broad monocyte-recruiting signature observed following genotoxic stress.

Consistent with this inflammatory phenotype, CD68 expression was also significantly increased following HG exposure (**Fig. 5v**). The presence of elevated CD68 alongside SASP cytokine induction suggests that metabolically stressed microglia adopt a persistently activated state, despite differences in downstream inflammatory signalling pathways compared with genotoxic stress.

Collectively, these findings demonstrate that both genotoxic and metabolic stress induce senescence-associated inflammatory responses in microglia, accompanied by sustained upregulation of the activation marker CD68. Dox-induced stress drives a robust, chemokine-rich SASP characterised by broad upregulation of canonical pro-inflammatory cytokines and monocyte-recruiting chemokines, consistent with a DNA damage-associated senescence programme. In contrast, HG-induced stress promotes a distinct inflammatory profile marked by induction of interferon-γ signalling and selective upregulation of CCL3, CCL4 and CCL5 in the absence of significant CCL2 induction, consistent with a metabolically driven stress response. These stimulus-specific SASP signatures, together with persistent CD68 expression, are likely to differentially shape chronic neuroinflammatory environments by sustaining microglial activation and immune signalling in the absence of cell loss.

## Discussion

In the present study, we demonstrate that chronic genotoxic and metabolic stress induce senescence-like phenotypes in human microglia that share core features but diverge in their regulatory pathways, mitochondrial adaptations and inflammatory outputs. These findings support the concept that cellular senescence represents a heterogeneous spectrum of states shaped by the initiating stressor rather than a uniform cellular endpoint [16,5,18].

Both Dox-induced genotoxic stress and HG-induced metabolic stress converged on key hallmarks of senescence, including cellular and nuclear hypertrophy, increased SA-β-gal activity, reduced metabolic activity and persistent DNA damage signalling. The activation of the p53-p21 axis under both conditions is consistent with a core mechanism of stress-induced growth arrest [20,6]. However, important differences emerged in cell-cycle regulation, with HG uniquely inducing p16 expression and promoting peripheral nuclear localisation of p21. These findings suggest that, despite convergence on growth arrest, distinct upstream stressors engage divergent regulatory programmes. The preferential activation of p16 under metabolic stress aligns with previous reports linking chronic metabolic perturbation to stabilised, long-term senescence states, whereas genotoxic stress primarily drives a p53-dependent response [18,19].

Importantly, control experiments using equimolar Mtl indicated that some responses, including p53, p21 and p16 induction, are sensitive to osmotic perturbation and may reflect a general stress response rather than glucose-specific signalling [28]. However, Mtl failed to reproduce key features of the HG condition, particularly NRF2 activation and coordinated KEAP1-NRF2-TFAM signalling, supporting the conclusion that metabolic stress engages distinct regulatory mechanisms beyond osmotic effects [26,29].

Mitochondrial dysfunction emerged as a central and shared feature of both stress paradigms, consistent with its established role in senescence and microglial ageing [21,30]. Both Dox and HG reduced mitochondrial mass, indicating impaired mitochondrial homeostasis. However, the adaptive responses diverged markedly. Under genotoxic stress, reduced NRF2 and TFAM expression suggests impaired activation of antioxidant defence and mitochondrial biogenesis pathways, indicative of a failure to mount an effective compensatory response. This is in line with reports that persistent DNA damage can suppress mitochondrial recovery mechanisms and reinforce senescence [31,32,30]. In contrast, HG induced transcriptional upregulation of KEAP1-NRF2-TFAM signalling, suggesting activation of redox-sensitive pathways. However, this response was not functionally effective, as evidenced by reduced HMOX1 protein expression, indicating an uncoupling between transcriptional activation and downstream antioxidant output. Such dysregulated NRF2 signalling has been reported in chronic metabolic stress and ageing contexts, where compensatory responses become maladaptive [33].

Another key distinction between the two stress conditions is apoptotic signalling. Dox-treated microglia exhibited increased procaspase-3 levels without evidence of cytochrome c release, indicating apoptotic priming without execution. This phenotype is characteristic of senescent cells, which often resist apoptosis despite activation of upstream pathways [34,35]. In contrast, HG-treated cells showed transcriptional upregulation of both CASP3 and the anti-apoptotic regulator BCL2, supporting a survival-oriented adaptation. The induction of BCL2 is particularly notable, as it has been implicated in the persistence of senescent cells and represents a key mechanism of apoptosis resistance [36]. Together, these findings suggest that both stressors promote survival of dysfunctional microglia, although through distinct mechanisms.

A central finding of this study is that the inflammatory output of senescent microglia is strongly shaped by the initiating stressor. Dox-induced senescence was characterised by a robust, chemokine-rich SASP, with broad upregulation of canonical pro-inflammatory cytokines (IL6, IL1A, IL1B, IL8) and chemokines (CCL2-CCL5). This profile is consistent with the classical DNA damage-associated SASP, which promotes immune cell recruitment and reinforces senescence through paracrine signalling [37,38]. In contrast, HG-induced senescence displayed a more heterogeneous SASP profile, with preserved cytokine induction but selective chemokine expression and a notable increase in IFNG. This suggests engagement of distinct immune-modulatory pathways under metabolic stress, potentially reflecting altered immune-metabolic signalling [14,15].

The absence of significant CCL2 upregulation under HG conditions is particularly notable, as CCL2 is a key mediator of monocyte recruitment. This indicates that metabolic stress may differentially regulate immune cell trafficking compared to genotoxic stress. Meanwhile, enrichment of CCL3-CCL5 suggests selective activation of chemokine networks associated with microglial activation and immune crosstalk. These observations reinforce the emerging concept that the SASP is highly context-dependent and varies according to cell type and stressor [18,39].

The consistent upregulation of CD68 under both conditions further supports the presence of a persistently activated microglial phenotype. In the context of ageing and neurodegeneration, such chronically activated, senescent microglia are thought to contribute to sustained neuroinflammation and neuronal dysfunction [40,41]. Rather than undergoing apoptosis or clearance, these cells persist and may amplify inflammatory signalling within the brain microenvironment.

These findings have important implications for understanding microglial contributions to neurodegenerative disease. Genotoxic stress appears to drive a strongly pro-inflammatory, chemokine-dominant SASP that may enhance immune cell recruitment and exacerbate tissue inflammation. In contrast, metabolic stress promotes a more complex and heterogeneous inflammatory profile, including interferon-associated signalling, which may differentially influence disease progression. Given the strong links between metabolic disorders and neurodegeneration, these findings highlight a potential role for metabolically driven senescence in shaping brain inflammation [14].

Taken together, these findings support the concept that genotoxic and metabolic stress drive distinct senescence trajectories in human microglia, characterised by divergent regulation of cell-cycle arrest pathways, mitochondrial responses and inflammatory programmes. A schematic overview of the proposed mechanisms underlying these stress-specific senescence phenotypes is presented in **Fig 6**.

**Fig 6.**
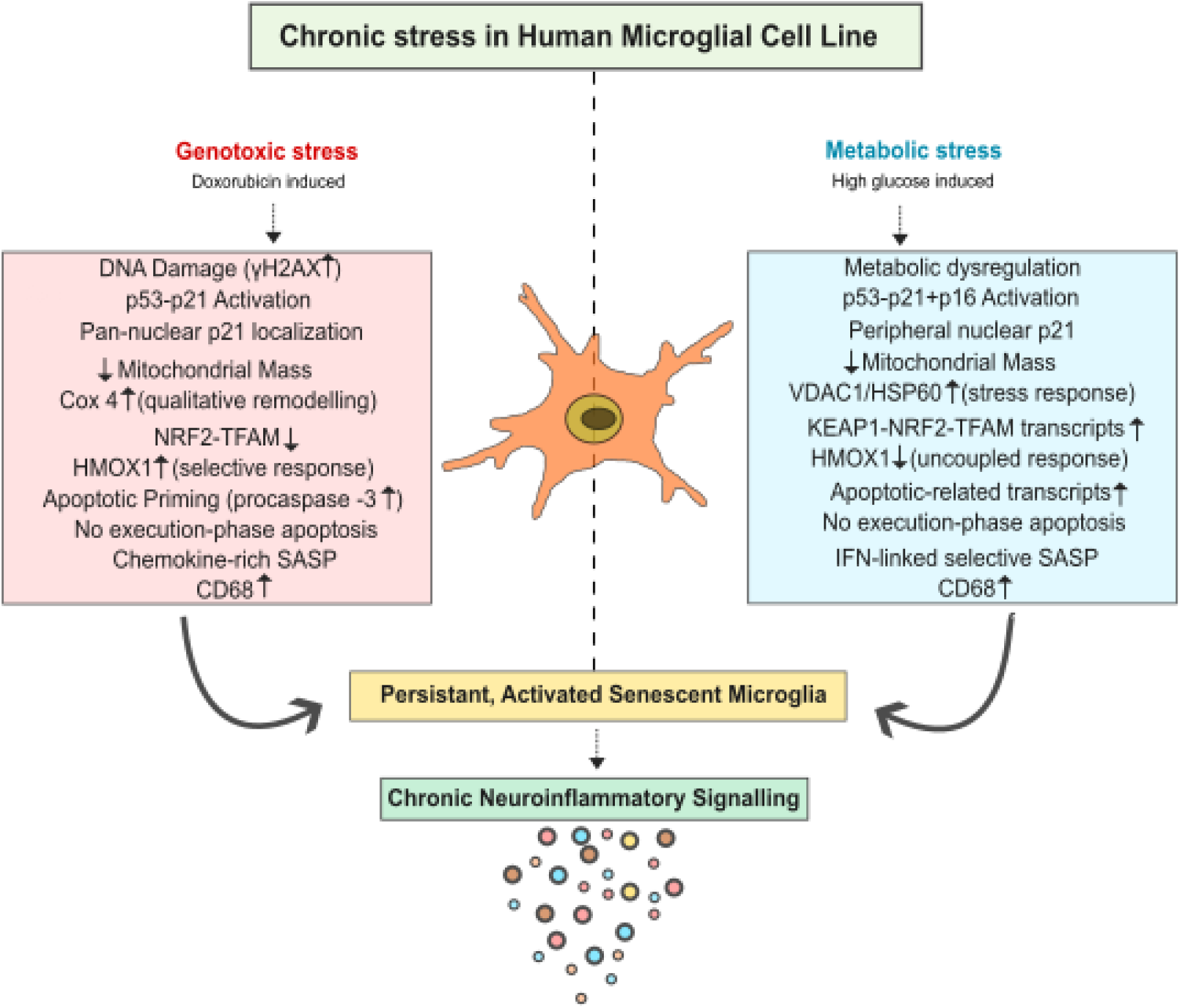
Schematic summary of doxorubicin– and high glucose-induced stress responses in human microglial cells. This schematic summarises the comparative cellular responses of human microglial cells to genotoxic stress (doxorubicin) and metabolic stress (high glucose). Doxorubicin induces DNA damage, evidenced by γH2AX formation and activation of the p53–p21 pathway, resulting in pan-nuclear localisation of p21, apoptotic priming (procaspase-3 accumulation) and the development of a chemokine-rich senescence-associated secretory phenotype (SASP), without progression to execution-phase apoptosis. In contrast, high glucose drives metabolic dysregulation characterised by activation of p53–p21 and p16, peripheral nuclear localisation of p21 and a distinct interferon-linked SASP profile. Although both stressors increase mitochondrial mass, doxorubicin is associated with qualitative mitochondrial remodelling (COX4), whereas high glucose induces a stress-response signature (VDAC1/HSP60). Furthermore, differential regulation of the KEAP1–NRF2–TFAM axis and HMOX1 expression indicates a selective or uncoupled antioxidant response between these conditions. Collectively, these pathways converge on a persistent, activated senescent microglial phenotype that sustains chronic neuroinflammatory signalling under prolonged stress.

Several limitations of this study should be acknowledged. The use of the immortalised HMC3 cell line, while enabling controlled mechanistic analysis, does not fully recapitulate the complexity of primary human microglia or the in vivo brain environment [23,42]. In addition, SASP pathway categorisation was based on predefined groupings rather than unbiased enrichment analysis. Future studies incorporating transcriptomic or proteomic approaches, as well as validation in primary microglia or in vivo models, will be important to confirm and extend these findings.

## Conclusion

In conclusion, our study demonstrates that chronic genotoxic and metabolic stress drive distinct yet overlapping senescence programmes in human microglia. These programmes are characterised by divergent cell-cycle regulation, mitochondrial adaptation and inflammatory signalling. The heterogeneity of these senescence states is likely to have significant functional consequences for microglial behaviour and the progression of neuroinflammatory and neurodegenerative diseases, highlighting the importance of stress-specific approaches in targeting senescent cells.

## Data Availability Statement

All data are available in the main text or the supplementary information. Raw data can be made available on reasonable request.

## Acknowledgements

We thank all members of the Platt and Kang labs for their support. We gratefully acknowledge the Microscopy and Histology Core Facility staff at the University of Aberdeen for their technical assistance.

## Author information

### Authors and Affiliations

Bettina Platt, Eunchai Kang, Nisha Vincy Jose, Susan Janssens: **Translational Neuroscience, Institute of Medical Sciences, University of Aberdeen, Scotland, United Kingdom.**

## Author Contributions

All authors (BP,EK,NVJ,SJ) contributed to the study conception, design, and data interpretation. Bettina Platt and Eunchai Kang supervised the study. Nisha Vincy Jose performed the laboratory work and data collection, drafted the first version of the manuscript and performed the data analysis. All authors (BP,EK,NVJ,SJ) contributed to reviewing and editing the manuscript and approved the final version.

## Corresponding author

Correspondence to Bettina Platt

## References

1. Hammond TR, Dufort C, Dissing-Olesen L, Giera S, Young A, Wysoker A, et al. Single-Cell RNA Sequencing of Microglia throughout the Mouse Lifespan and in the Injured Brain Reveals Complex Cell-State Changes. Immunity. 2019;50:253–271.e6. 10.1016/j.immuni.2018.11.004

2. Franceschi C, Garagnani P, Parini P, Giuliani C, Santoro A. Inflammaging: a new immune-metabolic viewpoint for age-related diseases. Nat Rev Endocrinol. 2018;14:576–90. 10.1038/s41574-018-0059-4

3. Leng F, Edison P. Neuroinflammation and microglial activation in Alzheimer disease: where do we go from here? Nat Rev Neurol. 2021;17:157–72. 10.1038/s41582-020-00435-y

4. Heneka MT, Carson MJ, El Khoury J, Landreth GE, Brosseron F, Feinstein DL, et al. Neuroinflammation in Alzheimer’s disease. Lancet Neurol. 2015;14:388–405. 10.1016/S1474-4422(15)70016-5

5. Gorgoulis V, Adams PD, Alimonti A, Bennett DC, Bischof O, Bishop C, et al. Cellular Senescence: Defining a Path Forward. Cell. 2019;179:813–27. 10.1016/j.cell.2019.10.005

6. Campisi J, d’Adda di Fagagna F. Cellular senescence: when bad things happen to good cells. Nat Rev Mol Cell Biol. 2007;8:729–40. 10.1038/nrm2233

7. Bhat R, Crowe EP, Bitto A, Moh M, Katsetos CD, Garcia FU, et al. Astrocyte senescence as a component of Alzheimer’s disease. PloS One. 2012;7:e45069. 10.1371/journal.pone.0045069

8. Wu J, Li M, Shi Y, Wang S. The senescent niche hypothesis: microglial dysfunction and replacement strategies in drug-resistant epilepsy. Front Immunol. 17:1807871. 10.3389/fimmu.2026.1807871

9. Streit WJ, Mrak RE, Griffin WST. Microglia and neuroinflammation: a pathological perspective. J Neuroinflammation. 2004;1:14. 10.1186/1742-2094-1-14

10. Antignano I, Liu Y, Offermann N, Capasso M. Aging microglia. Cell Mol Life Sci CMLS. 2023;80:126. 10.1007/s00018-023-04775-y

11. Hu Y, Fryatt GL, Ghorbani M, Obst J, Menassa DA, Martin-Estebane M, et al. Replicative senescence dictates the emergence of disease-associated microglia and contributes to Aβ pathology. Cell Rep. 2021;35:109228. 10.1016/j.celrep.2021.109228

12. Rolyan H, Scheffold A, Heinrich A, Begus-Nahrmann Y, Langkopf BH, Hölter SM, et al. Telomere shortening reduces Alzheimer’s disease amyloid pathology in mice. Brain J Neurol. 2011;134:2044–56. 10.1093/brain/awr133

13. Sedelnikova OA, Horikawa I, Zimonjic DB, Popescu NC, Bonner WM, Barrett JC. Senescing human cells and ageing mice accumulate DNA lesions with unrepairable double-strand breaks. Nat Cell Biol. 2004;6:168–70. 10.1038/ncb1095

14. Bernier L-P, York EM, MacVicar BA. Immunometabolism in the Brain: How Metabolism Shapes Microglial Function. Trends Neurosci. 2020;43:854–69. 10.1016/j.tins.2020.08.008

15. Lull ME, Block ML. Microglial activation and chronic neurodegeneration. Neurother J Am Soc Exp Neurother. 2010;7:354–65. 10.1016/j.nurt.2010.05.014

16. Torres G, Salladay-Perez IA, Dhingra A, Covarrubias AJ. Genetic origins, regulators, and biomarkers of cellular senescence. Trends Genet TIG. 2024;40:1018–31. 10.1016/j.tig.2024.08.007

17. Cohn RL, Gasek NS, Kuchel GA, Xu M. The heterogeneity of cellular senescence: insights at the single-cell level. Trends Cell Biol. 2023;33:9–17. 10.1016/j.tcb.2022.04.011

18. Hernandez-Segura A, Nehme J, Demaria M. Hallmarks of Cellular Senescence. Trends Cell Biol. 2018;28:436–53. 10.1016/j.tcb.2018.02.001

19. Wiley CD, Velarde MC, Lecot P, Liu S, Sarnoski EA, Freund A, et al. Mitochondrial Dysfunction Induces Senescence with a Distinct Secretory Phenotype. Cell Metab. 2016;23:303–14. 10.1016/j.cmet.2015.11.011

20. Kuilman T, Michaloglou C, Mooi WJ, Peeper DS. The essence of senescence. Genes Dev. 2010;24:2463–79. 10.1101/gad.1971610

21. Korolchuk VI, Miwa S, Carroll B, von Zglinicki T. Mitochondria in Cell Senescence: Is Mitophagy the Weakest Link? EBioMedicine. 2017;21:7–13. 10.1016/j.ebiom.2017.03.020

22. Masuda T, Sankowski R, Staszewski O, Prinz M. Microglia Heterogeneity in the Single-Cell Era. Cell Rep. 2020;30:1271–81. 10.1016/j.celrep.2020.01.010

23. Prinz M, Jung S, Priller J. Microglia Biology: One Century of Evolving Concepts. Cell. 2019;179:292–311. 10.1016/j.cell.2019.08.053

24. Dello Russo C, Cappoli N, Coletta I, Mezzogori D, Paciello F, Pozzoli G, et al. The human microglial HMC3 cell line: where do we stand? A systematic literature review. J Neuroinflammation. 2018;15:259. 10.1186/s12974-018-1288-0

25. Janabi N, Peudenier S, Héron B, Ng KH, Tardieu M. Establishment of human microglial cell lines after transfection of primary cultures of embryonic microglial cells with the SV40 large T antigen. Neurosci Lett. 1995;195:105–8. 10.1016/0304-3940(94)11792-h

26. Brownlee M. Biochemistry and molecular cell biology of diabetic complications. Nature. 2001;414:813–20. 10.1038/414813a

27. Nishikawa T, Edelstein D, Du XL, Yamagishi S, Matsumura T, Kaneda Y, et al. Normalizing mitochondrial superoxide production blocks three pathways of hyperglycaemic damage. Nature. 2000;404:787–90. 10.1038/35008121

28. Burg MB, Ferraris JD, Dmitrieva NI. Cellular response to hyperosmotic stresses. Physiol Rev. 2007;87:1441–74. 10.1152/physrev.00056.2006

29. Du C, Fang M, Li Y, Li L, Wang X. Smac, a mitochondrial protein that promotes cytochrome c-dependent caspase activation by eliminating IAP inhibition. Cell. 2000;102:33–42. 10.1016/s0092-8674(00)00008-8

30. Passos GS, Poyares D, Santana MG, Garbuio SA, Tufik S, Mello MT. Effect of acute physical exercise on patients with chronic primary insomnia. J Clin Sleep Med JCSM Off Publ Am Acad Sleep Med. 2010;6:270–5.

31. Correia-Melo C, Marques FDM, Anderson R, Hewitt G, Hewitt R, Cole J, et al. Mitochondria are required for pro-ageing features of the senescent phenotype. EMBO J. 2016;35:724–42. 10.15252/embj.201592862

32. Sahin E, Colla S, Liesa M, Moslehi J, Müller FL, Guo M, et al. Telomere dysfunction induces metabolic and mitochondrial compromise. Nature. 2011;470:359–65. 10.1038/nature09787

33. Bellezza I, Giambanco I, Minelli A, Donato R. Nrf2-Keap1 signaling in oxidative and reductive stress. Biochim Biophys Acta Mol Cell Res. 2018;1865:721–33. 10.1016/j.bbamcr.2018.02.010

34. Childs BG, Baker DJ, Kirkland JL, Campisi J, van Deursen JM. Senescence and apoptosis: dueling or complementary cell fates? EMBO Rep. 2014;15:1139–53. 10.15252/embr.201439245

35. Kirkland JL, Tchkonia T. Cellular Senescence: A Translational Perspective. EBioMedicine. 2017;21:21–8. 10.1016/j.ebiom.2017.04.013

36. Yosef R, Pilpel N, Tokarsky-Amiel R, Biran A, Ovadya Y, Cohen S, et al. Directed elimination of senescent cells by inhibition of BCL-W and BCL-XL. Nat Commun. 2016;7:11190. 10.1038/ncomms11190

37. Acosta JC, Banito A, Wuestefeld T, Georgilis A, Janich P, Morton JP, et al. A complex secretory program orchestrated by the inflammasome controls paracrine senescence. Nat Cell Biol. 2013;15:978–90. 10.1038/ncb2784

38. Coppé J-P, Patil CK, Rodier F, Sun Y, Muñoz DP, Goldstein J, et al. Senescence-associated secretory phenotypes reveal cell-nonautonomous functions of oncogenic RAS and the p53 tumor suppressor. PLoS Biol. 2008;6:2853–68. 10.1371/journal.pbio.0060301

39. Basisty N, Kale A, Jeon OH, Kuehnemann C, Payne T, Rao C, et al. A proteomic atlas of senescence-associated secretomes for aging biomarker development. PLoS Biol. 2020;18:e3000599. 10.1371/journal.pbio.3000599

40. Safaiyan S, Kannaiyan N, Snaidero N, Brioschi S, Biber K, Yona S, et al. Age-related myelin degradation burdens the clearance function of microglia during aging. Nat Neurosci. 2016;19:995–8. 10.1038/nn.4325

41. Heneka MT, Carson MJ, El Khoury J, Landreth GE, Brosseron F, Feinstein DL, et al. Neuroinflammation in Alzheimer’s disease. Lancet Neurol. 2015;14:388–405. 10.1016/S1474-4422(15)70016-5

42. Masuda T, Sankowski R, Staszewski O, Prinz M. Microglia Heterogeneity in the Single-Cell Era. Cell Rep. 2020;30:1271–81. 10.1016/j.celrep.2020.01.010

