## Supplementary Information for "Genotoxic and metabolic stress drive divergent senescence programs in human microglia" (from another version of this preprint: version 1, 2026-09-04)

**The PDF file includes:**

Supplementary Figure 1-3

Supporting data Figures 1 and 2

**Other Supplementary Information for this manuscript includes the following:**

Supplementary Tables 1, 2 and 3

Source Data

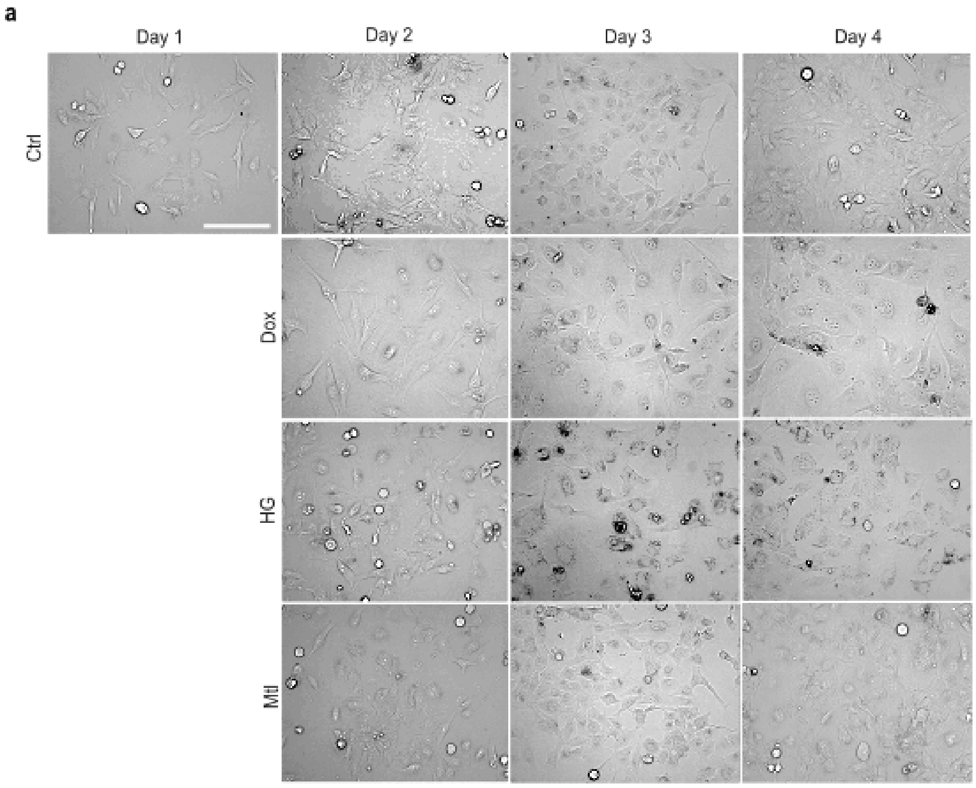


Jose et al. Supplementary Fig. 1

**Supplementary Fig. 1 Time-course images showing microglial morphology changes in culture**

(**a**) Representative brightfield time-course images showing microglial morphology in Ctrl, Dox, HG, and Mtl - treated HMC3 cells in culture. Scale bar: 100 µm

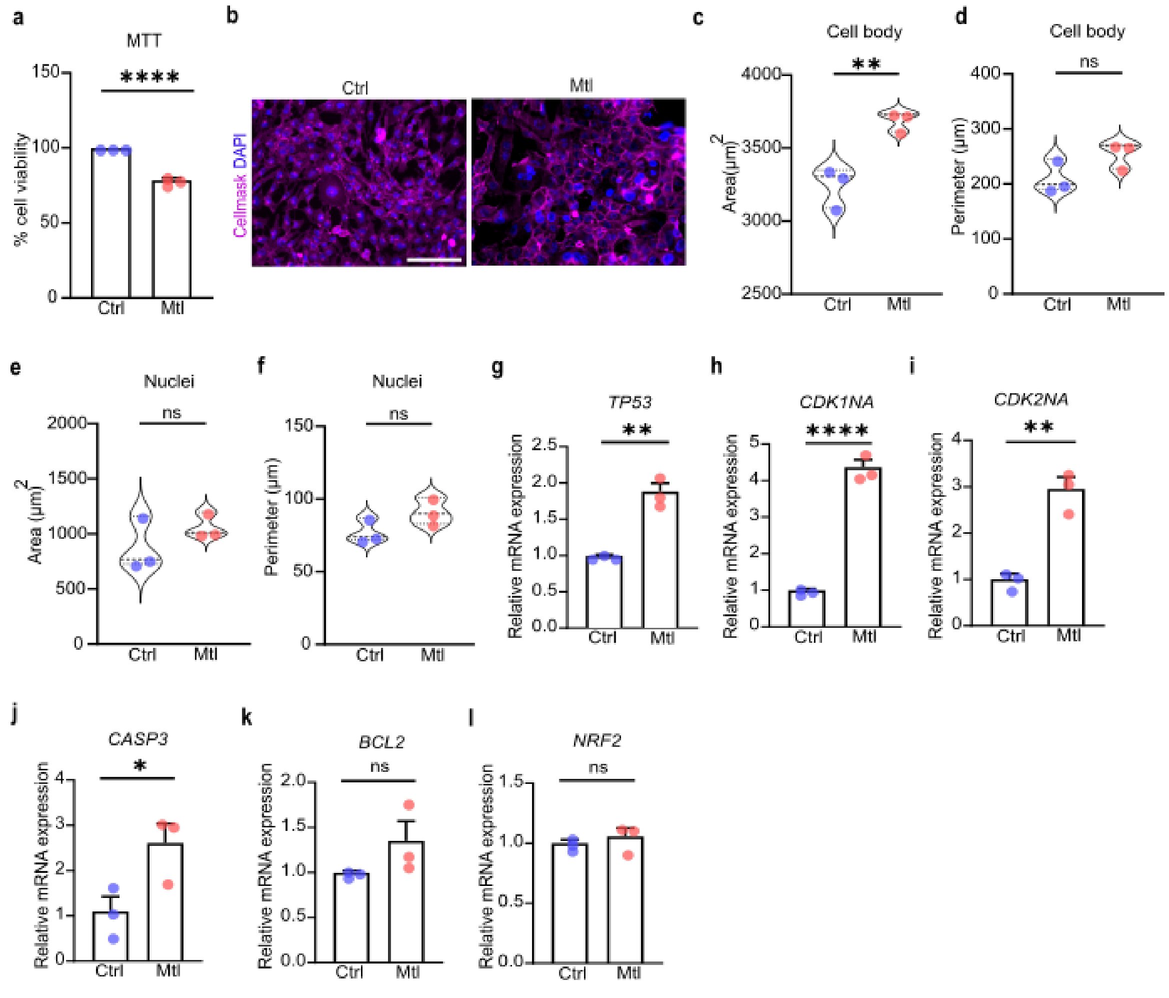


Jose et al. Supplementary Fig. 2

**Supplementary Fig. 2 Mannitol-induced osmotic stress effects on cell viability, morphology, activation of the p53-p21-p16 cell cycle arrest pathway, apoptosis, and NRF2 signaling.**

(**a**) MTT assay showing the effects of Mtl treatment on HMC3 cell viability. Values represent mean + SEM from $n = 3$ biological replicates, ****$p < 0.0001$, unpaired two-tailed Student's $t$ test.

(**b**) Representative images of morphology staining showing microglial morphology of Ctrl and Mtl-treated HMC3 cells. Scale bar: 100 µm

(**c-d**) Quantification of cell body areas and perimeters in Ctrl and Mtl-treated HMC3 cells. Values represent mean + SEM from $n = 5$ biological replicates, ns; not significant, **$p < 0.01$, unpaired two-tailed Student's $t$ test.

**(e-f)** Quantification of nuclear areas and perimeters in Ctrl and Mtl-treated HMC3 cells. Values represent mean + SEM from $n = 5$ biological replicates, ns; not significant, unpaired two-tailed Student's *t* test.

**(g–i)** Relative mRNA expression levels of TP53 (p53) (A), CDKN1A (p21) (B), and CDKN2A (p16) (C) in Ctrl and Mtl-treated-HMC3 cells measured by RT–qPCR. Values represent mean+ SEM from n = 3 biological replicates, normalized to the housekeeping genes RPS18 and GAPDH, $^{**}p < 0.01$; $^{****}p < 0.0001$, unpaired two-tailed Student's t test.

**(j-k)** Relative mRNA expression levels of CASP3 (D) and BCL2 (E) in Ctrl and Mtl-treated-HMC3 cells measured by RT–qPCR. Values represent mean+ SEM from n = 3 biological replicates, normalized to the housekeeping genes RPS18 and GAPDH, ns; not significant, $^{*}p < 0.05$, unpaired two-tailed Student's t test.

**(l)** Relative mRNA expression levels of NRF2 (F) in Ctrl and Mtl-treated-HMC3 cells measured by RT–qPCR. Values represent mean+ SEM from n = 3 biological replicates, normalized to the housekeeping genes RPS18 and GAPDH, ns; not significant, $^{*}p < 0.05$, unpaired two-tailed Student's t test.

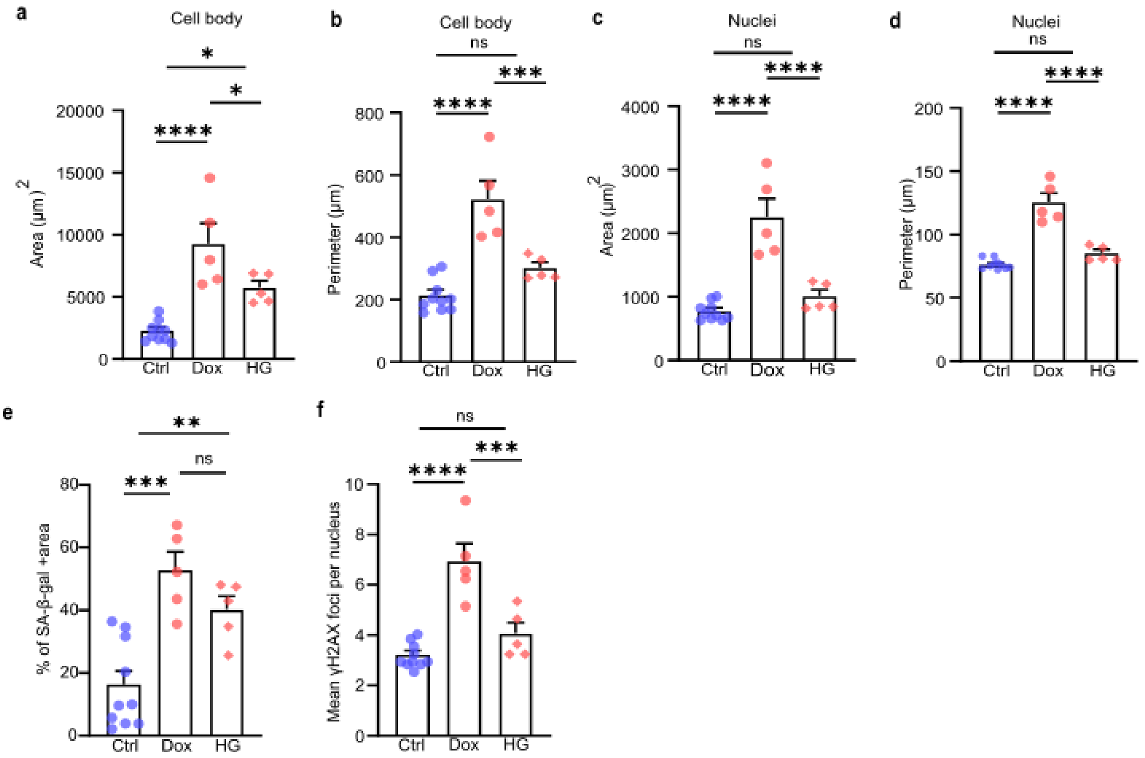


Jose et al. Supplementary Fig. 3

**Supplementary Fig. 3 Multiple comparisons analysis of senescence-associated morphological and DNA damage markers in HMC3 cells following genotoxic and metabolic stress**

**(a-b)** Quantification of cell body areas and perimeters in Ctrl, Dox and HG-treated HMC3 cells. Values obtained from independent experiments and are presented as mean + SEM, ns; not significant, **p* < 0.05; ****p* < 0.001; *****p* < 0.0001, ordinary one-way ANOVA followed by Tukey's multiple comparisons test.

**(c-d)** Quantification of nuclei areas and perimeters in Ctrl, Dox and HG-treated HMC3 cells. Values obtained from independent experiments and are presented as mean + SEM, ns; not significant, *****p* < 0.0001, ordinary one-way ANOVA followed by Tukey's multiple comparisons test.

**(e)** Quantification of the area percentage of senescence-associated β-galactosidase staining in Ctrl, Dox and HG-treated HMC3 cells. Values obtained from independent experiments and are presented as mean

+ SEM, ns; not significant, **$p < 0.01$; ***$p < 0.001$, ordinary one-way ANOVA followed by Tukey's multiple comparisons test.

**(e)** Quantification of the percentage of nuclei with γH2AX+ foci in Ctrl, Dox and HG-treated HMC3 cells. Values obtained from independent experiments and are presented as mean + SEM, ns; not significant, ***$p < 0.001$; ****$p < 0.0001$, ordinary one-way ANOVA followed by Tukey's multiple comparisons test.

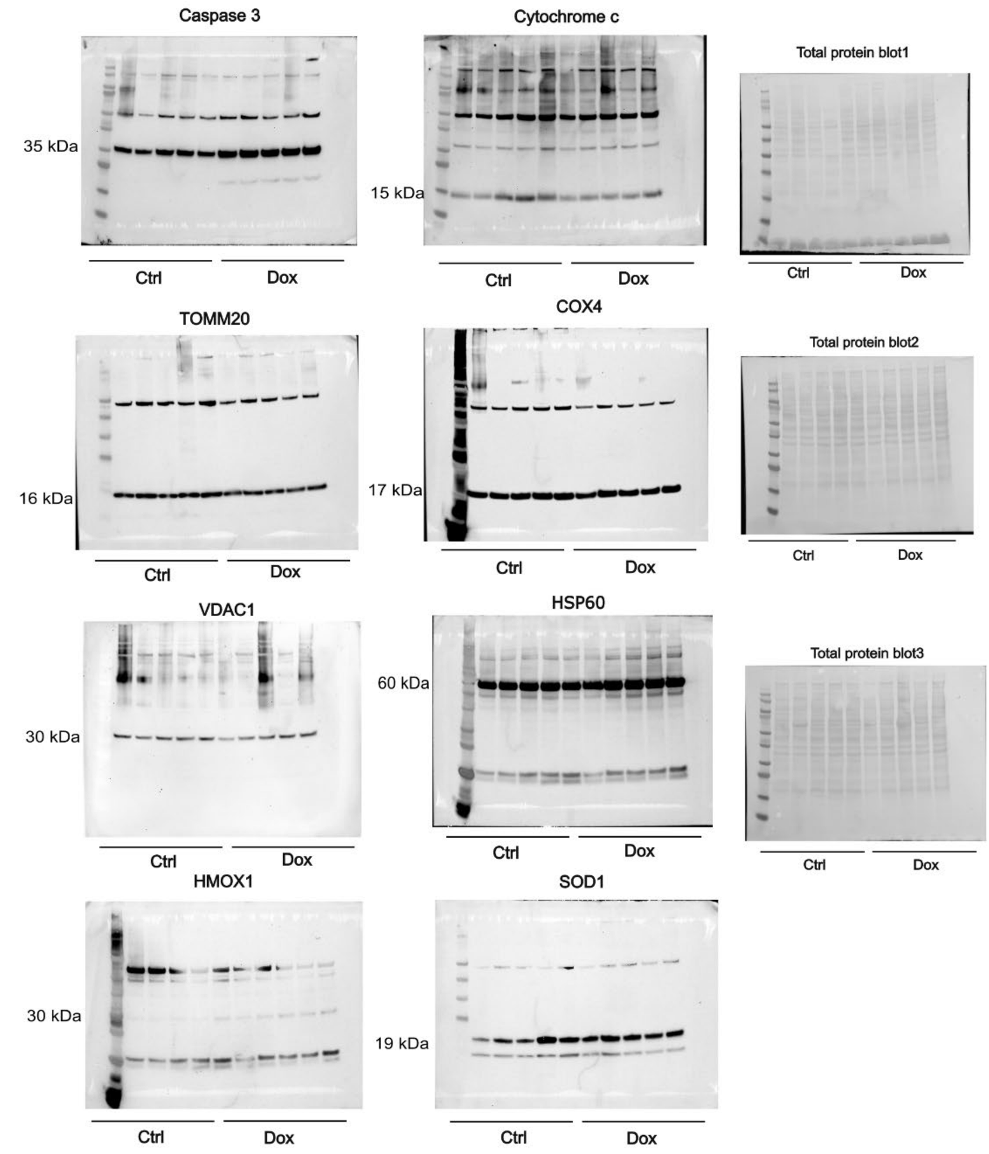


| Lane | Total protein |
|---|---|
| Ctrl 1 | 89221.7 |
| Ctrl 2 | 72737.4 |
| Ctrl 3 | 109662.3 |
| Ctrl 4 | 111286.6 |
| Ctrl 5 | 88199.3 |
| Dox 1 | 84101.7 |
| Dox 2 | 107149.2 |
| Dox 3 | 98504.8 |
| Dox 4 | 106522.7 |
| Dox 5 | 113856.7 |

| Lane | Total protein |
|---|---|
| Ctrl 1 | 78634.6 |
| Ctrl 2 | 65779.3 |
| Ctrl 3 | 101540.3 |
| Ctrl 4 | 119763.4 |
| Ctrl 5 | 95498.7 |
| Dox 1 | 33620.3 |
| Dox 2 | 60757.0 |
| Dox 3 | 64510.0 |
| Dox 4 | 55572.8 |
| Dox 5 | 86738.6 |

| Lane | Total protein |
|---|---|
| Ctrl 1 | 94876.8 |
| Ctrl 2 | 91878.1 |
| Ctrl 3 | 104377.9 |
| Ctrl 4 | 104179.3 |
| Ctrl 5 | 93561.3 |
| Dox 1 | 50075.5 |
| Dox 2 | 70064.5 |
| Dox 3 | 82141.3 |
| Dox 4 | 56651.6 |
| Dox 5 | 50271.8 |

**Supporting data Fig.1 Full-length uncropped Western blot images corresponding to main figures**

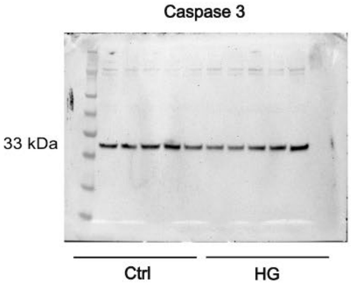


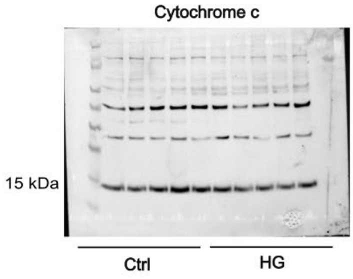


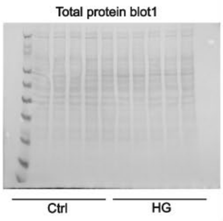


| Lane | Total protein |
|---|---|
| Ctrl 1 | 112955.5 |
| Ctrl 2 | 115718.5 |
| Ctrl 3 | 118566.1 |
| Ctrl 4 | 130208.0 |
| Ctrl 5 | 115027.7 |
| HG 1 | 95147.7 |
| HG 2 | 83613.9 |
| HG 3 | 73214.0 |
| HG 4 | 71668.6 |
| HG 5 | 92787.1 |

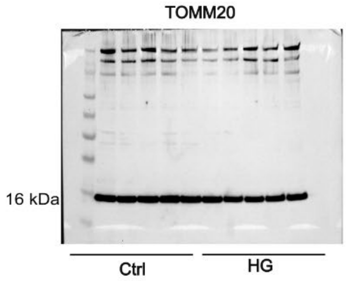


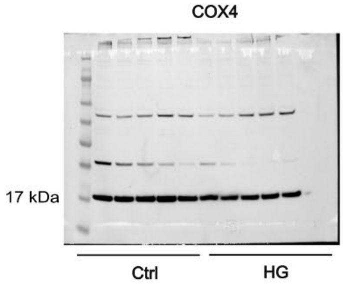


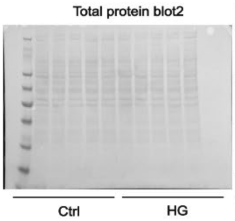


| Lane | Total protein |
|---|---|
| Ctrl 1 | 158722.7 |
| Ctrl 2 | 124885.8 |
| Ctrl 3 | 107805.8 |
| Ctrl 4 | 116541.5 |
| Ctrl 5 | 117742.9 |
| HG 1 | 118074.1 |
| HG 2 | 93544.4 |
| HG 3 | 79149.1 |
| HG 4 | 74641.8 |
| HG 5 | 76599.9 |

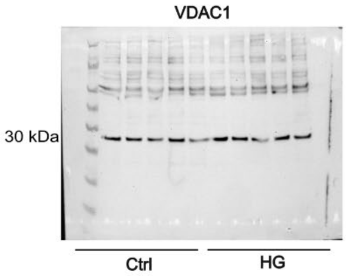


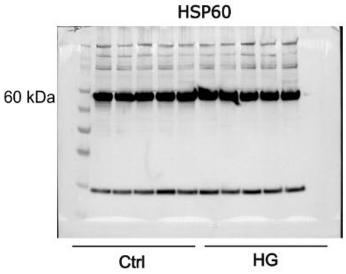


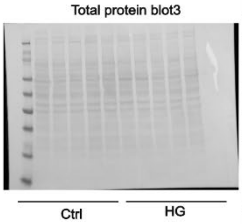


| Lane | Total protein |
|---|---|
| Ctrl 1 | 126237.7 |
| Ctrl 2 | 100950.8 |
| Ctrl 3 | 102636.5 |
| Ctrl 4 | 92001.2 |
| Ctrl 5 | 108869.3 |
| HG 1 | 98331.8 |
| HG 2 | 78460.2 |
| HG 3 | 68480.7 |
| HG 4 | 74362.6 |
| HG 5 | 78874.5 |

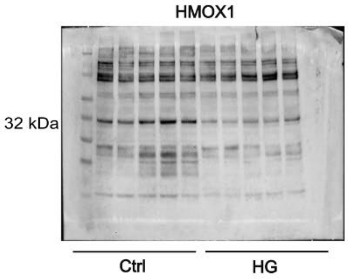


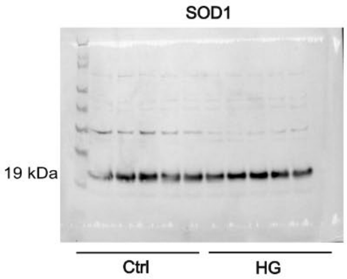


**Supporting data Fig.2 Full-length uncropped Western blot images corresponding to main figures**
